# Gene-specific endothelial programs drive AVM pathogenesis in SMAD4 and ALK1 loss-of-function

**DOI:** 10.1101/2025.01.03.631070

**Authors:** Olya Oppenheim, Wolfgang Giese, Hyojin Park, Elisabeth Baumann, Andranik Ivanov, Dieter Beule, Anne Eichmann, Holger Gerhardt

**Author notes:** Corresponding author: Prof. Holger Gerhardt. Co-corresponding author: Dr Olya Oppenheim Max-Delbrück-Center for Molecular Medicine in the Helmholtz Association (MDC) Robert-Rössle-Strasse 10 13125 Berlin Germany.

## Abstract

**Background:** Hereditary hemorrhagic telangiectasia is a genetic disorder caused by loss-of-function mutations in components of the bone morphogenetic protein signaling pathway, leading to arteriovenous malformations. Most prior work has treated BMP-component mutations as mechanistically interchangeable, yet whether distinct genes converge on a shared mechanism remains unclear. We aimed to understand the molecular relationship between BMP signaling and endothelial flow response that leads to AVM formation.

**Methods:** We expose human endothelial monolayers treated with siRNA against SMAD4 or ALK1 to laminar flow and analyze flow-responsive transcriptomics, flow-responsive BMP signaling activation dynamics, cell polarity and morphology. We analyze the cell-autonomous and non-cell-autonomous migration dynamics of ECs treated with siSMAD4 or siALK1. Using the postnatal mouse retina model, we study EC distribution changes over time in mosaic settings, and assess the remodelling capabilities of Smad4^iECKO^ or Alk1^iECKO^, relative to littermate controls.

**Results:** This study shows that mutations in SMAD4 or ALK1 lead to fundamentally distinct mechanisms of malformation formation. SMAD4 deficiency enhances endothelial responses to blood flow, including transcriptional activation and migration against flow, causing excessive capillary pruning and the development of single large shunts. In contrast, ALK1 deficiency disrupts flow sensing, impairs cell polarization and migration, and promotes a persistent angiogenic state, resulting in dense, hypervascularized networks. RNA sequencing reveals that these transcriptional changes precede flow onset, suggesting early defects in endothelial fate specification. Mosaic in vitro models show that mutant cells co-opt neighboring wild-type cells, while in vivo tracking confirms mutation-specific migration behavior.

**Conclusions:** These findings reveal divergent cellular programs driving arteriovenous malformations and underscore the need for gene-specific diagnostic and therapeutic strategies.

**Clinical Perspective:** *What is New?:*

- This study identifies gene-specific mechanisms of AVM formation in endothelial cells with SMAD4 or ALK1 loss-of-function, two major mutations underlying Hereditary Hemorrhagic Telangiectasia.
- Using multiple side-by-side analyses, the study reveals that SMAD4 deficiency enhances flow-mediated remodeling, while ALK1 deficiency impairs flow sensing and promotes angiogenic persistence.
- RNA sequencing showed that several transcriptional differences precede the onset of flow, indicating early divergence in endothelial cell fate specification and mechanosensory behavior.
- Mosaic culture and retinal models demonstrate that mutant cells exert distinct non-cell-autonomous effects on wild-type endothelial cells.

*What Are the Clinical Implications?:*

- This work challenges the prevailing assumption of a unified pathogenic pathway in HHT and supports a model of mutation-specific AVM pathogenesis, with distinct cellular and molecular drivers.
- The findings provide a mechanistic basis for the variable clinical presentation seen in patients with different HHT mutations.
- Identification of early transcriptional signatures, such as persistent ANGPT2 or DPP4 upregulation, may offer new biomarkers or therapeutic targets for early intervention before AVMs become clinically symptomatic.
- Personalized therapeutic strategies could reduce reliance on invasive treatments and shift toward mechanism-based, precision medicine approaches in HHT and related vascular disorders.

## Introduction

The vascular system develops and remodels through a series of complex cellular processes that continuously adapt its shape and function to the local needs of tissues and organs^1–6^. Defects in the fine-tuned coordination of endothelial cell (EC) collective migration, proliferation, perfusion-mediated stabilization and cellular rearrangements in response to fluid shear stress (FSS) can lead to vascular malformations that can severely threaten organ function, or cause bleedings and life-threatening complications. Understanding the precise molecular and cellular mechanisms that drive aberrant vessel formation is fundamental to the development of precision therapy.

Central to such understanding is the behavior of ECs during formation and remodeling of the vascular network. Exactly how the collective behavior of ECs leads to the formation of a vascular network with hierarchical organization of arteries, capillaries and veins remains incompletely understood. Whereas the initial vasculogenic assembly of vessels^2,5^ and subsequent sprouting angiogenesis^3^ are largely independent of blood flow, the stabilization of nascent vessels and the orderly pruning process that leads to the mature network depend on both the biomechanical forces of blood flow and related biochemical signaling events^1,7–14^. Recent work identified flow-migration coupling as the orderly migratory response of ECs to changing FSS which drives functional remodeling. Failure of ECs to engage in this flow-migration coupling process has been shown to occur in animal models of Hereditary Hemorrhagic Telangiectasia (HHT)^7,9,10,15–18^.

HHT is an autosomal dominant vascular disorder^19^, and is characterized by development of arteriovenous malformations (AVMs). Such malformations can occur in many organs and can be small (dilated small blood vessels called telangiectasias, typically appearing on the skin and nasal-mouth mucosa) or large intraorgan AVMs^19,20^.

The major genetic effectors causing this disorder are part of the Bone Morphogenetic Protein (BMP) signaling pathway^21–24^. Loss-of-function mutations in the co-receptor Endoglin^25^, the type I receptor ALK1 and downstream DNA-binding protein SMAD4 account for the majority of diagnosed cases, and lead to HHT1^25^, HHT2^26^ or JP/HHT^22^, respectively.

When circulating BMP ligands bind to the extracellular domain of the ALK1-ENG heterotetramer, a conformational change in the intracellular domains subsequently leads to phosphorylation of receptor SMADs 1/5/9. These phosphorylated SMADs then bind to SMAD4 and translocate to the nucleus, where this complex regulates target-gene expression. As homozygous loss-of-function mutations in any of the effectors is embryonic lethal, accumulation of somatic mutations in the healthy allele drives the loss-of-function phenotype, in a process called loss of heterozygosity^23^. Shunts appear in a sporadic manner and do not affect the entire vascular network within a tissue. Moreover, different somatic mutations accumulate in different cells and vascular beds^27^. For these shunts to start developing in a mature and quiescent network, a “second hit” vascular injury, which causes a reactivation of the vascular bed and the exit from a quiescent state, is usually required to occur^27,28^. Current treatment options include mostly symptomatic and surgically invasive approaches, while mechanism-based drug options are limited^19,29^.

The unpredictable behavior, variable presentation, and limited therapeutic options of AVMs pose significant clinical challenges, underscoring the need to elucidate the underlying mechanisms. In our search for critical and common molecular mediators and cellular mechanisms of HHT, we find that loss of SMAD4 or ALK1 surprisingly shows very different effects and influences the EC response to FSS in opposite ways, ultimately leading to different AVM pathomechanisms. Loss of SMAD4 enhances collective migration against flow, leading to hyperpruning of proximal capillaries, therefore precipitating a shunt. In contrast, ALK1 deficient cells show a delayed and reduced migration against flow, coupled with angiogenic persistence, causing AVMs to grow via hypopruning and enlargement of capillaries. These findings provide valuable insights into distinct pathomechanisms that lead to AVM formation, offering potential avenues for improved diagnostics and mechanism based treatment options.

## Methods

### Cell culture experiments

Human umbilical venous endothelial cells (HUVECs, PromoCell) were expanded and used in passages 2-4. Culture flasks were coated with 0.2% gelatin prior to seeding. Cells were cultured in EGM2-Bulletkit (Lonza) or MV2 (PromoCell) in a 37°C humidity incubator with 5% CO_2_. For siRNA knockdown, cells were seeded in T25 flasks and transfected after 24h at 60-70% confluence for *siCTRL* (Qiagen AllStars negative control, 40pmol), *siSMAD4* (Qiagen, FlexiTube GeneSolution for SMAD4: SI03089527, SI03042508, SI00076041 and SI00076020; 10pmol each) or *siALK1* (Qiagen, FlexiTube GeneSolution for ALK1: SI02659972 and SI02758392, 10pmol each), using RNAiMax transfection reagent (Thermofisher) diluted in OptiMem media and added to antibiotic free EBM2 media (Lonza). After 5h, transfection mix was aspirated and replaced with full media.

### End point flow experiments

HUVECs were harvested 24h after siRNA knockdown and seeded onto gelatinized 0.4 Luer ibiTreat slides (Ibidi) at a concentration of 2 million cells per ml and placed overnight in a CO_2_ incubator. The following day, slides were connected to perfusion units (Ibidi) and exposed to 0.6 Pa or 1.8 Pa shear stress for 4h. Cells that were exposed to shear stress for 16h would be connected later on, and would have a media exchange earlier that day. A static control for each condition was placed in the flow incubator for the same duration as the flow experiments. After each flow experiment, cells would either be fixated with 4% PFA for subsequent immunofluorescence assays, or lysed with RLT buffer (RNAeasy RNA extraction kit, Qiagen) for RNA based subsequent analysis.

### RNA extraction for qPCR, RNA seq

After flow experiments, cells were lysed with two rounds of 200ul RLT buffer and collected into tubes. Cell lysates were kept in -80°C freezers until all relevant samples have been collected, then RNA extraction would be completed with the RNAeasy kit according to manufacturer’s instructions, including DNAse step. RNA concentration was assessed using NanoDrop. RNA was sent to the genomic core facility for RNA sequencing, or used for qPCR analysis. cDNA synthesis was carried out using BioRad’s or Thermofisher’s cDNA kits according to manufacturer’s instructions. qPCR was performed using Taqman probes on a QuantStudio 6 qPCR machine in 384 well plates.

### Bioinformatic analysis

RNA-Seq reads were mapped to the human genome (GRCh38.p7) using STAR (version 2.7.3a^30^). The reads were assigned to genes with FeatureCounts (version 2.0.3^31^) using Gencode version 25 (Ensembl 85) annotation and the following parameters “ -t exon -g gene_id -s 2 -p”.

Differential expression analysis was carried out using DESeq2 (version 1.38^32^) using default parameters. Only genes with at least 5 reads in at least 3 samples were considered for the analysis. Gene set enrichment analysis was carried out with CERNO test from the R tmod package (version 0.50.13^33^) using MsigDB Biological Pathways GO collection.

### Immunofluorescence, imaging of *in vitro* flow samples

After fixation, slides were blocked and stained with primary antibodies against VEcadherin (goat, AF938, R&D Systems, 1:1000); GM130 (mouse, 610822, BD Bioscience, 1:500); pSMAD159 (rabbit, 13820S, Cell Signaling, 1:500) or KLF4 (rabbit, HPA002926, Sigma Aldrich, 1:500). Secondary antibodies were used in 1:400 dilution: Donkey anti-goat IgG 568 (A11077, Thermofisher); Donkey anti-mouse IgG 488 (A21202, Thermofisher); Donkey anti-rabbit IgG 647 (A31573, Thermofisher). Nuclei were stained using a 1:1000 DAPI solution. Mowiol mounting media was mixed with Dabco solution (Sigma) at 10:1 ratio and applied to slides. Slides were imaged on the Zeiss 980 Confocal inverted microscope with a 20x air objective by taking 10 non overlapping z-stack images per slide.

### Image analysis of *in vitro* data

Images were processed using sum intensity projection and were then analyzed using the Polarity-JaM toolbox and web-application^34^. The full code is available at https://github.com/polarityjam. Polarity analysis (for nuclei-Golgi polarity, cell orientation) was performed via acquisition of a polarity index (PI), which is an indicator of the concentration of the circular distribution, its value (limited between 0 and 1) indicating the collective orientation strength of the monolayer^35^. In addition, the signed polarity index (V-score) indicates the strength of polarization with respect to a given direction. The signed polarity index varies between -1 and 1 and indicates the strength of polarization with respect to an assumed direction of polarization.

Additional features were extracted through the web application www.polarityjam.com:

Nuclear marker accumulation was analyzed via the ratio of the mean intensity of pSMAD159/KLF4 in the nucleus and cytosol (NUC/Cyt ratio). Cell area was extracted and normalized to µm^2^. Cell elongation was extracted as the major/minor axis ratio. Circular statistics was performed via the PolarityJaM web application. Cell orientation of each condition was extracted as V-score per image from the circular statistics file.

### Live migration under flow

For live migration experiments, cells were prepared and seeded in the same conditions as described for the end-point flow experiments. On the day of the experiment, seeding MV2 media was aspirated and replaced with CO2 independent media (Promocell basal media without phenol-red and without sodium bicarbonate, supplemented with MV2 supplement kit, B-glycerolphosphate at a final concentration of 4.32ug/ul and sodium bicarbonate at a final concentration of 0.0075%) with addition of Spy505 (Spyrochrome) nuclear dye at 1:1000 dilution and incubated at 37°C for 4h. Fluidic units would be placed inside the incubation chamber of the microscope and warmed up prior to connection of slides. Slides were connected and placed on multi slide stage insert. Time Lapse imaging would run for 48h and acquire 3×3 tiles with 3 slice Z-stacks for each position, with a time interval of 7.5 minutes. Migration analysis would then be done using the TrackMate feature^36^ on Fiji (ImageJ) with StarDist segmentation tool^37^. Tracks would then be analyzed using a Python based script^38^ on Jupyter notebook (AnaConda).

For mosaic migration experiments, prior to seeding into flow slides, cells were labeled with CellTracker Green or Red (Thermofisher) with 1:5000 dilution (final concentration 2nM) in EBM2 media and incubated for 45 minutes in 37°C, then washed three times with full MV2 media and placed in the incubator for about 4h. Cells were then trypsinized and harvested, and seeded in the same concentration with 1:1 ratio between green and red cells. Live imaging would begin the following day, 48h after siRNA treatment. Time Lapse imaging was conFig.d in the same manner, and ran for approx. 2.5 days (500 cycles with 7.5 minute intervals). Subsequent analysis for each channel was done in the same manner as described in the previous paragraph.

### *In vivo* experiments

All animal experiments were performed under a protocol approved by the Institutional Animal Care Use Committee of Yale University (no.2023-11406).

Seven to eight weeks old ALK1^f/f^ or SMAD4 ^f/f^ mice with Cdh5 CreERT2 and mTmG mixed genetic background were intercrossed for experiments and ALK1^f/f^ Cdh5 CreERT2 (mTmG) mice or SMAD4 ^f/f^ Cdh5 CreERT2 (mTmG) were used. Gene deletion was induced by intra-gastric injections with 100 μg Tamoxifen (Sigma, T5648; 2.5 mg ml−1) into pups at P4 or P1-3 to stain Collagen type 4 and CD31, and 0.75 μg Tamoxifen into P5 pups for mosaic knockout and labeling. Tx-injected CreERT2 negative littermates were used as controls. Retinae were collected on P6 for Collagen type 4 and CD31 staining, and on P8 and P15 for mosaic population analysis.

### Immunostaining and imaging of mouse retinae

Retinae were prefixed in 4% PFA for 8 min at room temperature. Retinae were dissected, blocked for 30 min at room temperature in blocking buffer (1% fetal bovine serum, 3% BSA, 0.5% Triton X-100, 0.01% Na deoxycholate, 0.02% Sodium Azide in PBS at pH 7.4) and then incubated with specific antibodies in blocking buffer overnight at 4C. The next day, retinae were washed and incubated with IB4 together with the corresponding secondary antibody overnight at 4°C, then washed and post-fixed with 0.1% PFA and mounted in fluorescent mounting medium (DAKO, USA). High-resolution pictures were acquired using a Leica SP8 confocal microscope with a Leica spectral detection system (Leica TCS SP8 detector), and the Leica application suite advanced fluorescence software. Primary antibodies: IB4 ([IsolectinB4] #132450, 1:400; Life Technologies), GFP Polyclonal Antibody, Alexa Fluor 488 (#A-21311, 1:1000; Invitrogen), CD31 (553370; 1:200; BD), Collagen IV antibody (2150-1470; 1:400; Bio-Rad).Secondary antibodies: Alexa Fluor 488 anti-Rat (A21208, 1:500, Invitrogen), Alexa Fluor 568 anti-Rabbit (A10042, 1:500, Invitrogen).

### Image analysis of *in vivo* data

Analysis of mosaic labeled mouse retinae was done as previously described^39,40^. Briefly, maximum projections of the IB4 channel were used to create drawn masks of retina outline, veins, arteries and optic nerve in Fiji^41^. Maximum projections of the GFP channel were used to create GFP masks. For every pixel in the GFP mask, three numbers were computed (using drawn masks as referential): (1) distance to the nearest vein (d_v_); (2) distance to the nearest artery (d_a_); and (3) radial distance to the optic nerve (d_r_). From these measures, the relative distances by ϕ_v-a_ = d_v_/(d_v_ + da) were obtained. The EC distribution was computed by performing the operation for x randomly selected GFP-positive pixels in each retina, which were used as a proxy for EC distribution. A kernel density estimation was used to approximate the underlying EC distribution in the two-dimensional coordinate system spanned by ϕ_v-a_ and d_r_. For computational analysis, a Python-based workflow was used, accessible on GitHub https://github.com/gerhardt-lab/retina-VACS-HHT

Analysis of mouse retinae used in regression analysis was done using the 3DVascNet software^42^. Briefly, z-stacks of CD31 and ColIV channels were uploaded into the software, along with a resolution file depicting the micron/pixel ratio of each image. Channels of the same retina were segmented independently from each other. After segmentation, a region of interest (ROI) was selected for each image, generally encompassing one retina leaflet. ROIs are skeletonized in 3D using the generated 3D mask from the segmentation step. The software outputs feature quantifications, including ROI volume and number of branching points. Regression events for each retina are quantified using a composite of the CD31 and ColIV channels and within the borders of the analyzed ROI. A regression event is classified by an absence of the CD31 signal, while ColIV staining remains intact. Partial absence of CD31 indicates the vessel is in an intermediate regression stage; complete absence of CD31 while ColIV staining remains intact indicates the final regression stage of a vessel; complete absence of CD31 and partial disruption of ColIV staining indicates long regressed vasculature. Regression frequency is quantified via the proportion of regression events per 100 branching points in the CD31 channel. Branching points and Vessel density ratios are quantified by dividing the outputs of the CD31 by ColIV channels, and subsequent normalization to the ROI volume ratio.

### Statistical Analysis - DABEST method

In order to compute effect sizes along with common p-value statistics, we used the DABEST (’data analysis with bootstrap-coupled estimation’) method^43^. The mean difference distributions including 95% confidence intervals were plotted to indicate effect size. Common statistical analysis include two-sided Welch’s test, student t-Test and a non-parametric Mann-Whitney test. The test used is indicated in the Fig. captions.

### Use of large language models (LLMs)

Large language models (LLMs), including ChatGPT (OpenAI), were used to assist with non-scientific aspects of manuscript preparation, including rephrasing existing text for clarity and readability. In addition, LLMs were used to support the adaptation of existing Python scripts for statistical data analysis, where package updates or deprecated commands required modification. No novel analyses were generated by the model, and all scientific interpretations and data validation were performed by the authors. The updated scripts were added to Github at https://github.com/gerhardt-lab/retina-VACS-HHT.

## Results

### RNA sequencing unveils distinct transcriptional regulation associated with ALK1 and SMAD4 loss-of-function

In search for common pathogenic downstream effectors of ALK1 and SMAD4 loss-of-function we performed RNA sequencing on flow-mediated shear-stress exposed HUVECs following SMAD4 or ALK1 siRNA mediated knock-down (supplementary Fig. 1.1). The FSS magnitude was adjusted to 0.6 Pa to simulate FSS values typically occurring in veins and capillaries, as previous work has shown that AVMs in HHT models commence in capillaries and veins, but not in arteries^21^. RNA samples were collected after 4 and 16 hours of flow exposure (Fig. 1a). Principal component analysis (PCA) of all samples showed samples clustered primarily according to flow duration (Fig. 1b), with untreated controls (light gray dots) and control siRNA treated samples (dark gray dots, *siCTRL* hereafter) clustering together. Surprisingly however, ALK1 siRNA and SMAD4 siRNA-treated samples did not cluster together. This is remarkable, given that SMAD4 is a downstream component of the BMP9/10 signaling pathway triggered by the activation of ALK1. Instead, *siALK1*-treated samples (magenta dots) clustered separately from *siSMAD4*-treated samples (blue dots), and samples treated with both *siSMAD4* and *siALK1* (siDouble hereafter, green dots), clustered between the two single treatments in all time points. We therefore set out to investigate the differences in gene regulation in more detail.

**Figure 1:**
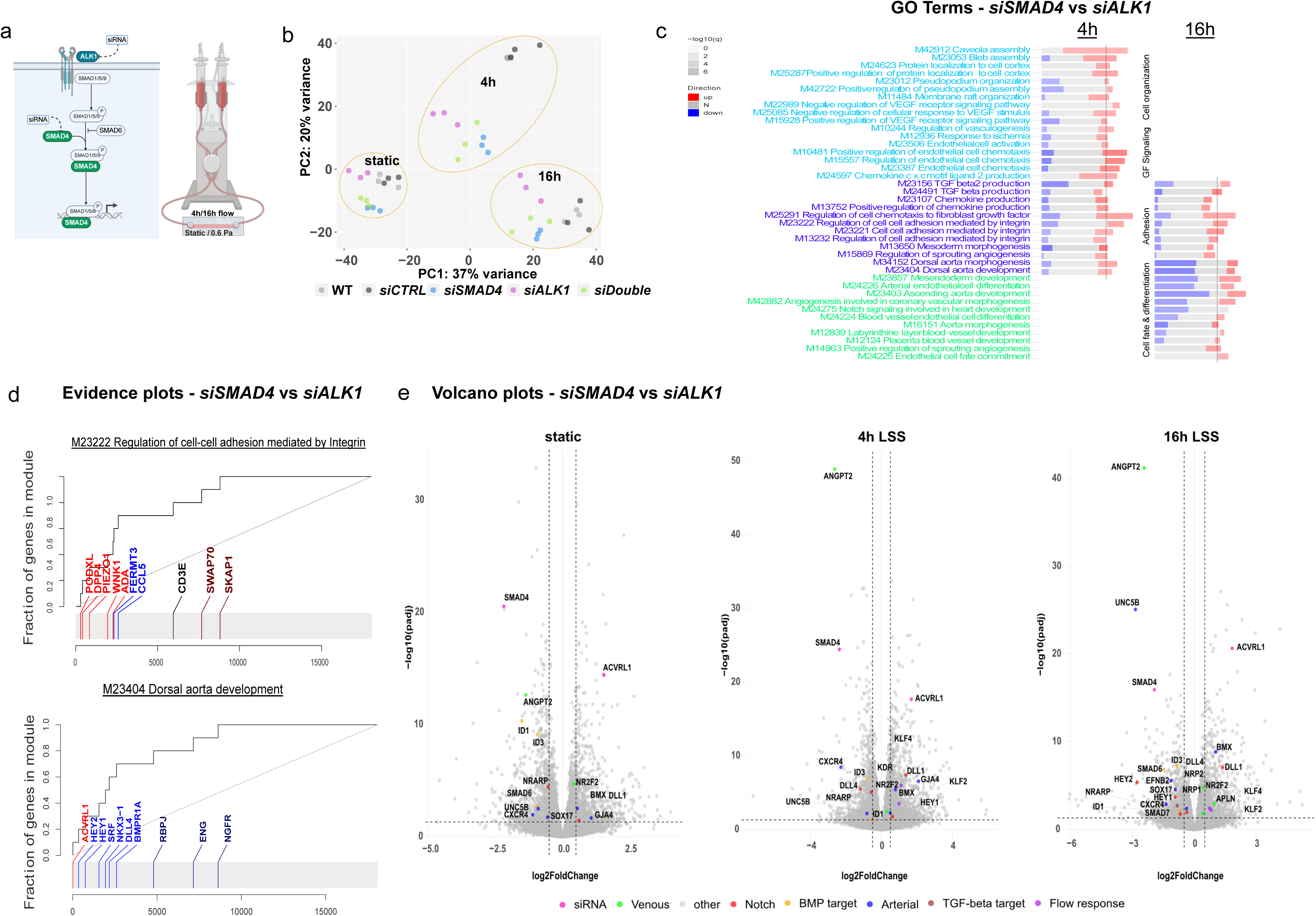
differential transcriptomic changes in ECs under flow. a. Experimental schematic-HUVECs treated with siRNA against ALK1 or SMAD4 are exposed to 4 or 16 hours of laminar FSS 0.6 Pa (LSS hereafter). b. Principal component analysis of RNAseq samples. Dashed circles group samples by flow duration. n=3 independent experiments. c. Panel plot of select biological processes, significantly (adj. P-value<0.05) enriched in *siSMAD4*-treated samples vs *siALK1*-treated samples after 4 and/or 16h of LSS. The length of the rectangle is the effect size (area under the curve or AUC, see Figure 1d). In red and blue are the fraction of significantly up- or down-regulated genes from that gene set. AUC_min_=0, AUC_max_=1. Vertical lines represent threshold value (AUC=0.65) for each column. d. Evidence plots of GO Terms representing differential regulation in two biological processes. X-axis is the list of all genes sorted by their p-value. Y-axis is the cumulative fraction. Light blue and light red colors represent significant down– or up-regulation, respectively. e. Volcano plots of differentially expressed genes in *siSMAD4* treated samples vs *siALK1* treated samples, significance cut-off -Log2FC of at least 0.5. Select genes of interest are color-annotated. SMAD4 and ACVRL1 are annotated in pink to confirm knockdown.

Gene set enrichment analysis using GO biological processes (GO_BP hereafter) directly comparing *siSMAD4*-treated samples versus *siALK1*-treated samples revealed 6 clusters, based on flow dependence and duration (supplementary Fig. 1.2). Flow-dependent GO_BP (Fig. 1c) were divided into 3 clusters - differential expression at early flow exposure (light blue), flow-dependent differential expression (dark blue), and differential expression at late flow exposure (light green). Interestingly, the early and late flow response GO_BP were quite different, with early flow exposure showing an enrichment in upregulation of biological processes associated with cell compartment organization, chemotaxis, VEGF and TGF-beta signaling in *siSMAD4*-treated samples, whereas late flow exposure exhibited enrichment in downregulation of GO_BPs associated with cell fate and arterial differentiation in *siSMAD4* treated samples, compared to *siALK1* treated samples.

Evidence plots and volcano plots from the early flow response as well as late flow response exposed many differentially expressed genes, suggesting ALK1 and SMAD4 loss-of-function leads to highly differential effects on the endothelial response to flow (Fig. 1d-e and supplementary Fig. 1.3).

### ALK1 is crucial for flow-induced BMP pathway activation, while SMAD4 governs its duration

As SMAD4 is a direct transcriptional regulator of the ALK1-activated BMP pathway but also the TGF-beta pathway, it was essential to understand the roles of both ALK1 and SMAD4 on flow-induced BMP pathway activation. To test this, we exposed HUVECs to low shear stress of 0.6 Pa (LSS hereinafter) for 4 and 16 hours as in the RNAseq setup and assessed BMP pathway activation via the intensity of the nuclear pSMAD159 signal (Fig. 2 a, b). In static conditions, HUVECs displayed basal nuclear pSMAD159 levels. Exposing HUVECs to 4h of LSS showed increased and variable activation of the BMP pathway, which however subsided after 16h in the *siCTRL*-treated samples. The same flow regime in *siSMAD4*-treated samples led to a more uniform activation of the BMP pathway that remained active at 16h, while in *siALK1*-treated samples no flow-induced activation of the BMP pathway was observed. Similar outputs were obtained for high shear stress levels (1.8 Pa, HSS hereinafter, Fig. 2c and supplementary Fig. 2). Taken together, these data suggest an essential role for ALK1 in flow-induced activation of the BMP pathway, while SMAD4 absence results in a prolonged, more robust and uniform BMP pathway activation.

**Figure 2:**
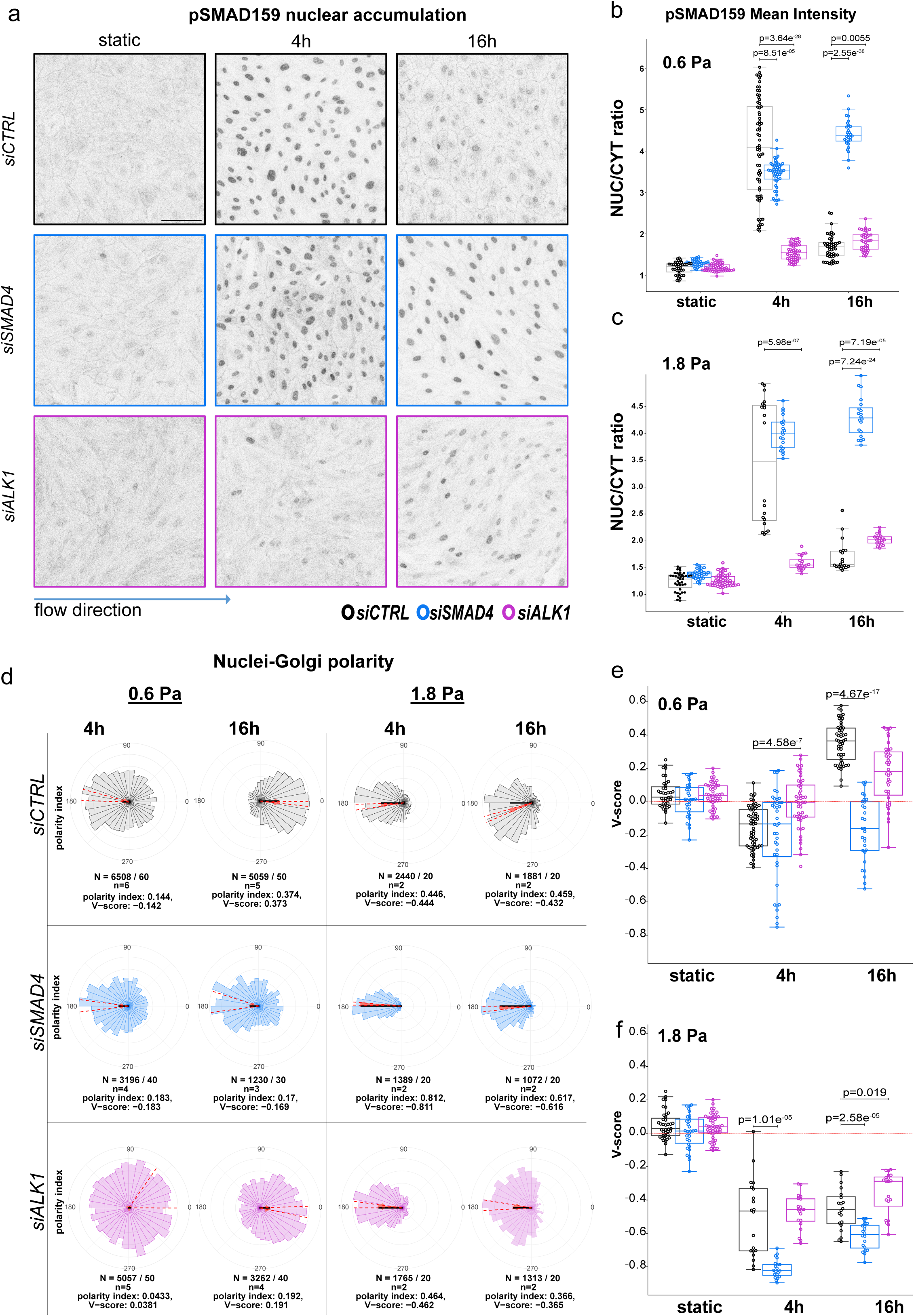
BMP pathway activation and cell polarity under flow. a. pSMAD159 staining of HUVECs after exposure to LSS. Control cells have an initial activation of the BMP pathway which goes down over time, while *siSMAD4* cells have constant BMP activity and siALK1 cells don’t have BMP pathway activation. Scale bar 100μm. b. and c. Quantification of nuclear pSMAD159 signal, normalized to cytosolic signal at 0.6 and 1.8 Pa. Statistical analysis was performed using Welch’s test, n≥30 images per condition, from at least 3 independent experiments (b) and n=20 from 2 independent experiments (c). d. Rose diagrams depicting the Nuclei-Golgi polarity of *siCTRL-*, *siSMAD4-* and *siALK1-*treated ECs exposed to LSS or HSS (1.8 Pa) for 4 and 16 hours. The red arrow points towards the mean direction of polarization and its length represents the polarity index (PI), which is an indicator for the variance of the distribution; the black line represents the signed polarity index (V-score). Data underneath each rose diagram depict the total number of cells/images (N), number of experiments (n) and numerical values for PI and V-Score. e. and f. Graphical representation of V-score values from d for LSS and HSS (f), each data point representing mean V-Score of an individual image. Box plots display median values along with the standard deviation. Statistical analysis was performed using Welch’s test.

### SMAD4 and ALK1 deficiency differentially impact cell polarity in endothelial cells

Cell polarity is a crucial aspect of development that is often characterized by the asymmetric distribution of organelles in response to extracellular stimuli such as biophysical forces^44^. ECs exhibit a striking ability to establish flow-directed polarity, characterized by the positioning of the Golgi apparatus against the direction of blood flow, during their migratory response^7,9,44^. There is growing evidence for changes in this front-rear polarity in AVMs, as well as in ECs depleted of BMP pathway components^21,24^. In order to characterize these changes, we analyzed the nuclei-Golgi polarity of HUVECs after exposure to LSS and HSS for 4 and 16 hours (Fig. 2d-f). In LSS conditions, we observed a weak polarization of *siCTRL*-treated samples against flow direction at 4h which reversed and intensified at 16h (Fig. 2 d, e). Surprisingly, *siSMAD4*-treated samples polarized against flow direction in both time points, whereas *siALK1*-treated samples exhibited random polarization at 4h, followed by a weak polarization with flow at 16h. When exposed to HSS, cells polarized against flow in all treatments although with strongest effects in *siSMAD4*-treated samples (Fig. 2 d, f). The fact that even *siALK1*-treated samples polarize against flow, although they do not show induction of pSMAD1/5/9, suggests that polarization induced by HSS occurs independent of BMP pathway activation. In summary, our data shows that ECs lacking SMAD4 or ALK1 respond very differently to FSS, particularly under LSS levels.

### FSS-dependent changes in cellular morphology are enhanced in the absence of SMAD4

A key function of flow-migration coupling of ECs is the fine tuning of vessel diameter. This fine tuning depends on the ability of ECs to accurately sense flow and to undergo cellular rearrangements that affect their morphology, size and orientation. Our data demonstrate that *siALK1*-treated samples fail to upregulate BMP signaling and to polarize against flow at LSS levels, suggesting they might have either lost the ability to sense flow, or to respond to flow. The expression of Krüppel-like 4 (KLF4) transcription factor has been established as flow sensitive^11,45,46^, as its mRNA levels increase in ECs upon exposure to flow. Analysis of KLF4 expression as surrogate for flow sensitivity confirmed that ECs sense the exposure to FSS under the different siRNA knockdown conditions. *siCTRL*- and *siSMAD4*-treated samples showed elevated KLF4 expression upon LSS exposure, and also *siALK1*-treated samples showed elevated KLF4 expression, albeit to a significantly lower magnitude (Fig. 3a). The notion that lack of ALK1 or SMAD4 differentially affects transcriptional responses to flow is further supported by the evidence plot of the biological process “Response to fluid shear stress” (Fig. 3b). Immunofluorescence staining for KLF4 protein also confirmed increased protein accumulation in the nucleus upon exposure to LSS. Albeit not to the same degree as seen at RNA level, *siSMAD4*-treated samples had elevated nuclear KLF4 accumulation in both time points (Fig. 3c, supplementary Fig. 3a). In addition, we stained ECs for the adherens junction marker VE-Cadherin, allowing us to segment and analyze cell area, elongation and orientation and evaluate which of these are flow-dependent and/or dependent on the loss of ALK1 or SMAD4 (Fig. 3d-g, supplementary Fig. 3b-e). Already in static conditions and throughout the flow experiment, *siALK1*-treated ECs in the monolayer formed distinct swirl-like clusters, where each sub-population maintained unique orientations independent of neighboring clusters (Supplementary Fig. 3b). After 16h of LSS, the cell area of *siALK1*-treated ECs remained similar to *siCTRL*-treated ECs (Fig. 3f, supplementary Fig. 3c). Although they were significantly more elongated (Fig. 3g), the elongation of *siALK1*-treated ECs was flow independent (Supplementary Fig. 3d). In contrast, *siSMAD4*-treated ECs had a significantly larger cell area compared to *siCTRL*- and *siALK1*-treated ECs (Fig. 3f, supplementary Fig. 3c). In addition, *siSMAD4*-treated ECs elongated and oriented themselves in a flow dependent manner (Fig. 3e,g and supplementary Fig. 3d,e). Taken together, these observations further suggest an enhanced sensitivity to LSS in absence of SMAD4 and reduced sensitivity to LSS in case of ALK1 absence.

**Figure 3:**
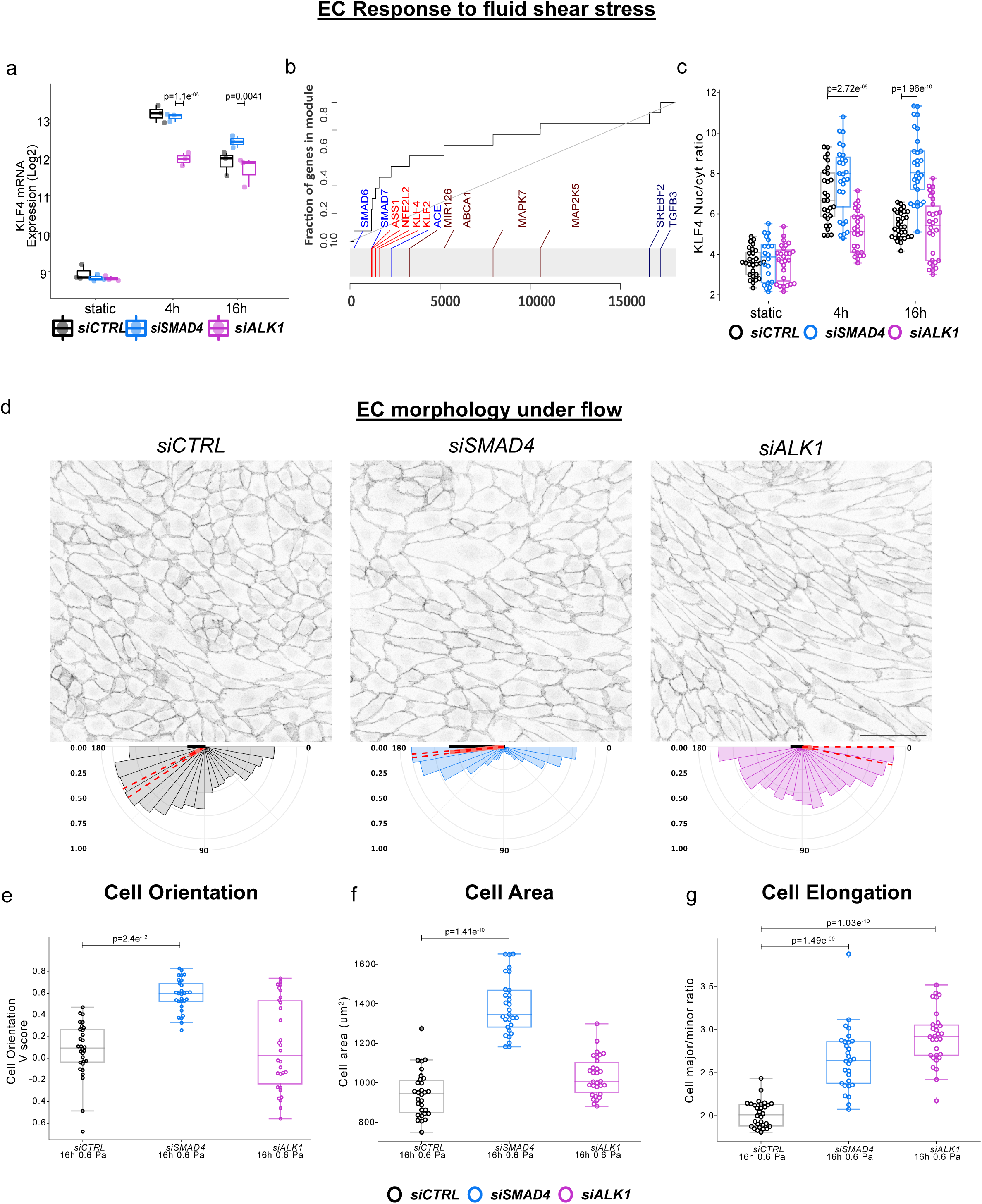
Endothelial response to FSS via KLF4 signaling and morphological changes. a. Analysis of KLF4 mRNA expression after exposure to LSS in Log2FC scale. Static samples are relative to wild-type static, flow samples are relative to static samples for the same siRNA condition. *siSMAD4*-treated samples have higher upregulation of KLF4 while *siALK1*-treated samples have a lower upregulation of KLF4, indicative of their disturbed flow sensation. Adjusted p-values (padj) are indicated. n=3 independent experiments. b. Evidence plot of biological process “response to fluid shear stress”, comparing *siSMAD4-* treated samples relative to *siALK1*-treated samples after 16h of LSS. X-axis is the list of all genes sorted by their p-value. Y-axis is the cumulative fraction. Light blue and light red colors represent significant down– or up-regulation. c. Quantification of nuclear KLF4 signal, normalized to cytosolic signal. Statistical analysis was performed using the Welch’s test, n=40 images per condition, from 4 independent experiments. d. Representative immunofluorescence staining of VEcadherin after 16h of LSS (top) and the corresponding cell orientation plots (bottom). Scale bar 100μm. e. Quantification of cell orientation V-score values. Box blots display median values and standard deviation. *siSMAD4*-treated cells orient themselves better parallel to flow than *siCTRL*-treated cells, and *siALK1*-treated cells form subpopulations that have different cell orientation. Statistical analysis was performed using Welch’s test, n=4 independent experiments with 10 images per experiment. f. Cell area analysis in μm^2^ and g. Cell elongation as ratio of major over minor axes of ECs after 16h of LSS. *siSMAD4*-treated cells are significantly bigger than *siCTRL*-treated cells; *siALK1*-treated cells are not bigger than *siCTRL*-treated cells but are less variable in size and shape. Statistical analysis using Welch’s test, n=4 independent experiments with 10 images per experiment.

### Endothelial cells exhibit distinct migration patterns in the absence of ALK1 or SMAD4

The end point analyses of SMAD4 and ALK1 deficient ECs so far captured two distinct phenotypes that suggest hypersensitivity and hyposensitivity to LSS, respectively. We therefore hypothesized that *siSMAD4*-treated ECs would migrate faster against flow, while *siALK1*-treated ECs would migrate less effectively against flow. By labeling the nuclei of ECs and tracking their movement over time under LSS conditions, we were able to analyze the migration pattern of ECs under different siRNA-treatment conditions (Fig. 4a-c, supplementary movies 1a-c). Analysis of individual cell tracks as well as population behaviour revealed that *siCTRL*-treated ECs exhibit a mild preference to migrate against flow (Fig. 4a, b left panels, red tracks), with a smaller proportion of cells migrating with flow (blue tracks) or perpendicular to flow. *siSMAD4*-treated ECs exhibited a much more prominent shift in trajectories against flow direction and also less perpendicular trajectories. The fraction of cells migrating with flow was reduced, but surprisingly, they moved with higher velocities (Fig. 4a, b middle panels). *siALK1*-treated ECs also showed an increased velocity as well as overall cell movement, but with the majority of cells migrating with flow. (Fig. 4a, b right panels). Plotting the average migration of all cells parallel to flow for each condition, revealed that *siCTRL*-treated ECs initially migrate against flow, then reverse direction to move with flow, and ultimately slow down to cease migrating after 48h. *siSMAD4*-treated ECs show a higher acceleration in net migration velocity against flow, but continue to migrate against flow throughout the duration of the experiment. *siALK1*-treated ECs initially behave similar to *siCTRL*-treated ECs, but rapidly decrease net migration velocity parallel to flow (Fig. 4c top panel).

**Figure 4:**
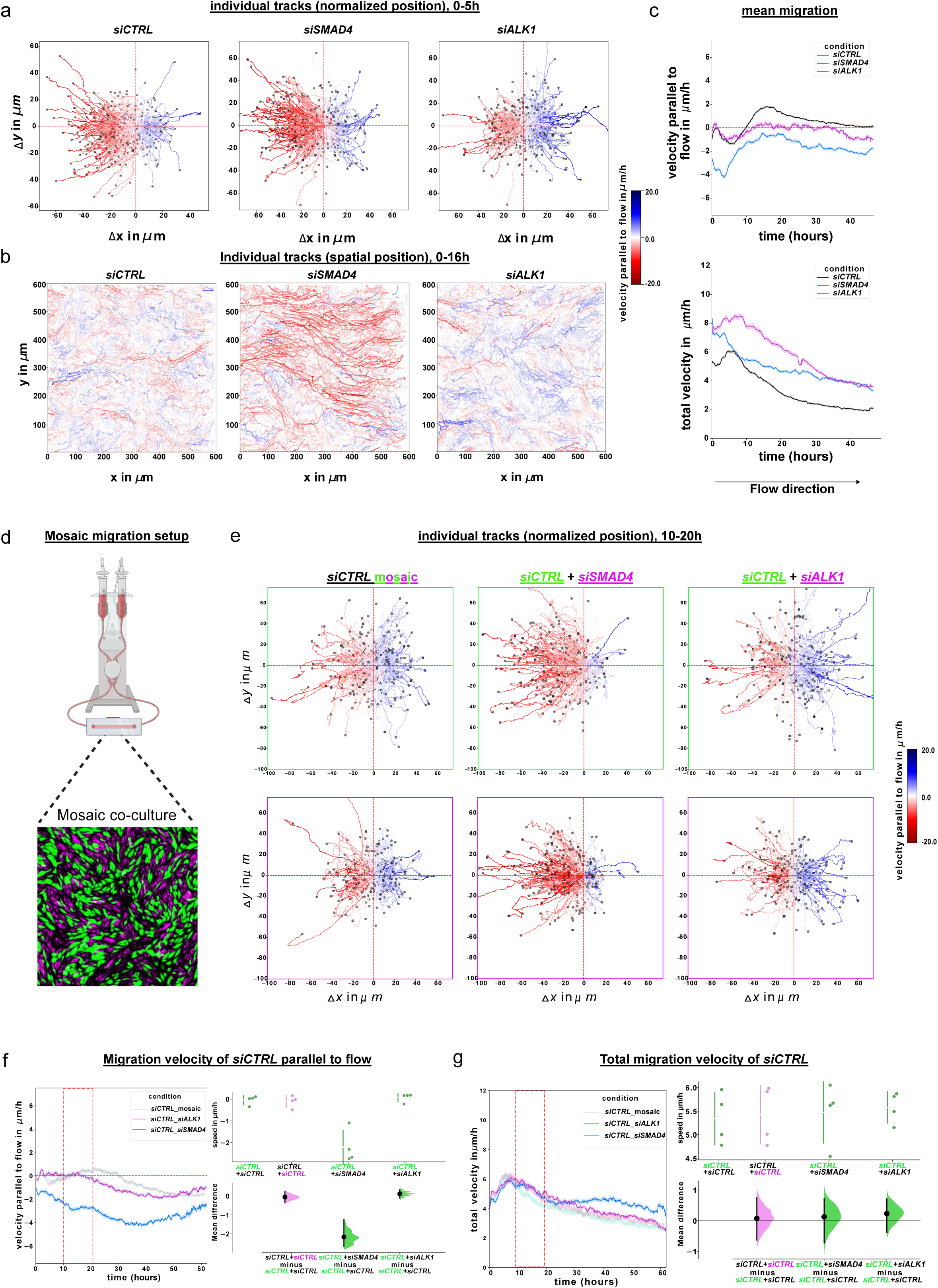
Endothelial migration *in vitro*. a. Bootstrapped trajectory plots of initial 5 hours of migration for all conditions, 100 tracks per replicate, and 400 tracks per condition. Starting point of each cell trajectory has been normalized to start from (0, 0). Color intensity map represents scale of parallel migration velocity with flow (blue) to against flow (red). b. Representative track images from individual samples of the initial 16 hours of migration. Spatial position of each track is kept. c. Mean migration velocity of *siCTRL-*, *siSMAD4-* or *siALK1*-treated HUVECs, parallel to flow direction (top) or overall (bottom). n=4 individual experiments per siRNA treatment. d. Mosaic migration setup. Cells are separately labeled with CellTracker Green (green cells) or Red (magenta cells) and mixed before seeding in flow slides. e. Bootstrapped trajectory plots with normalized starting position of the green (top) and magenta (bottom) cells in each co-culture, from t=10 to t=20 hours. 100 tracks per replicate per color, from n=4 independent experiments. Color intensity map represents scale of migration velocity with flow (blue) to against flow (red). f. and g. Mean migration velocity of *siCTRL*-treated cells when co-cultured either with themselves (*siCTRL*_mosaic, light pink/green curve), or with *siSMAD4*- or *siALK1*-treated cells (*siCTRL_siSMAD4*, *siCTRL_siALK1*, respectively), parallel to flow direction (f) or overall (g). Cells were labeled with CellTracker Green or Red (Thermofisher) and seeded together in 1:1 ratio. n=4 independent experiments. Red rectangles represent the time window of trajectory plot analysis. Statistical analysis: mean difference plots of the velocity parallel to flow (f) or total velocity (g), generated using estimation statistics. Both groups of *siCTRL*-treated cells (green+red) migrate similarly when cultured together. *siCTRL*-treated cells (green) migrate at a similar speed when cultured with *siALK1*-treated cells, or faster parallel to flow when cultured with *siSMAD4*-treated cells within the measured time interval (red dashed rectangles).

Intriguingly, *siCTRL*-treated cells show an average total velocity (irrespective of direction) that is slower than that of cells of the other conditions. *siALK1*-treated ECs show the highest initial velocity and slow down over time to match the speed of *siSMAD4*-treated ECs, but unlike the latter, they fail to migrate directionally against flow. These data strengthen the hypothesis that loss of SMAD4 renders ECs hypersensitivity to flow as they migrate more effectively against flow, while the loss of ALK1 leaves ECs unable to collectively respond to shear stress as they fail to migrate against flow, despite their high migratory potential. *siCTRL*-treated ECs display a very characteristic biphasic response and strong flow adaptation, suggesting that the rearrangements of ECs as a consequence of shear stress represent a transitory adaptation mechanism.

As HHT progresses by the loss of heterozygosity via somatic mutations on the healthy alleles of BMP pathway components in a subset of endothelial cells within a vascular plexus, a model for cell migration that more closely resembles the *in vivo* situation of HHT would be a mosaic scenario where only a subset of cells is deficient for SMAD4 or ALK1. By separately labeling the two populations, we generated and observed these mosaic scenarios under flow (Fig. 4d-g, supplementary movies 2a-c). Intriguingly, in the mosaic situation, when *siSMAD4*-treated cells were cultured together with *siCTRL*-treated cells, the latter group migrated more effectively against flow (Fig. 4e-middle panels and 4f-blue curve), initially without elevation of their overall average velocity, following the migration pattern of *siSMAD4*-treated cells; Interestingly, the overall average velocity of *siCTRL*-treated cells increased later on (Fig. 4g-blue curve). The opposite was observed when *siCTRL*-treated cells were cultured with *siALK1*-treated cells, as *siCTRL*-treated cells exhibited a noticeable shift towards migrating with the direction of flow, resulting in an average migration velocity of 0µm/h (Fig. 4e right panel and 4f-pink curve). Interestingly, this mosaic condition overcame the initial delay and displayed elevated migration against flow starting from the second day, preceding the *siCTRL*-mosaic condition by 10h. Taken together, these findings suggest that the absence of SMAD4 not only results in cell-autonomous hypersensitivity to LSS, but also that these cells affect control cells to migrate in a similar fashion. Likewise, the absence of ALK1 results in a cell-autonomous hyposensitivity of ECs to FSS, and also a non-cell autonomous effect on the co-migration of control cells in the mosaic situation.

### Cells lacking Alk1 or Smad4 exhibit distinct population movements in the retina *in vivo*

Should the loss of ALK1 or SMAD4 affect EC migration patterns in a similar way *in vivo*, we would predict that ALK1 knock-out ECs show a mild inability to migrate against flow, whereas SMAD4 knock-out cells migrate even more effectively than control cells. To assess this possibility, we turned to the mouse retina model, and employed mosaic endothelial Cre-lox mediated genetic labelling to follow population movements over time. Using our bespoke dual coordinate EC distribution analysis^39,40^, we tracked the changes in EC population positions in mosaic retinae overtime. CDH5-Cre^ERT2^ mTmG (CTRL^mTmG^ hereinafter), CDH5-Cre^ERT2^ Smad4^fl/fl^ mTmG (Smad4^mTmG^ hereinafter) or CDH5-Cre^ERT2^ Acvrl1^fl/fl^ mTmG (Alk1^mTmG^ hereinafter) pups were treated with a low dose (0.75ug) of Tamoxifen at P5, to induce mosaic knockout of Smad4 or Alk1, respectively (Fig. 5a). Retinae were collected at P8 and P15 and stained for GFP and IB4 (Fig. 5a-b). Using this approach, we have previously shown that EC populations shift from the vein and plexus to the artery, as cells migrate against the direction of flow^39,40^. The GFP+ EC density within the artery and surrounding arterioles was slightly reduced for Smad4^mTmG^ and Alk1^mTmG^ retinae at P8 relative to CTRL^mTmG^ (supplementary Fig. 4b-d – left; Fig. 5c – P8 KDE plots). Between P8 and P15 CTRL^mTmG^ retinae showed a prominent shift of cells to the artery (position 1.0 in the KDE plots at Fig. 5c, Fig. 5d – gray plots). However, this was even more pronounced in the Smad4^mTmG^ retinae (Fig. 5c-d, blue plots), but much less so in Alk1^mTmG^ retinae (Fig. 5c-d, pink plots). Further analysis of GFP+ ECs on the vein-artery axis confirmed a significant shift out of the peri-venous capillary bed for Smad4^mTmG^ as well as a shift into the peri-arterial capillary bed (Fig. 5e-f, blue plots). Alk1^mTmG^ retinae exhibit a weak shift within the peri-venous capillary bed, as well as a mild shift towards the peri-arterial capillaries and arteries, but insignificantly relative to control and much less than Smad4 mutant cells (supplementary Fig. 4c, pink plots, Fig. 5e-f, pink plots). Overall, these results demonstrate that mosaic loss of Smad4 or Alk1 in ECs *in vivo* leads to differential migration defects, in line with our observations in the *in vitro* flow assays. Whereas the absence of Smad4 enhances directional migration towards arterial structures, the absence of Alk1 seems to impair this directional EC migration, but not completely prevent it.

**Figure 5:**
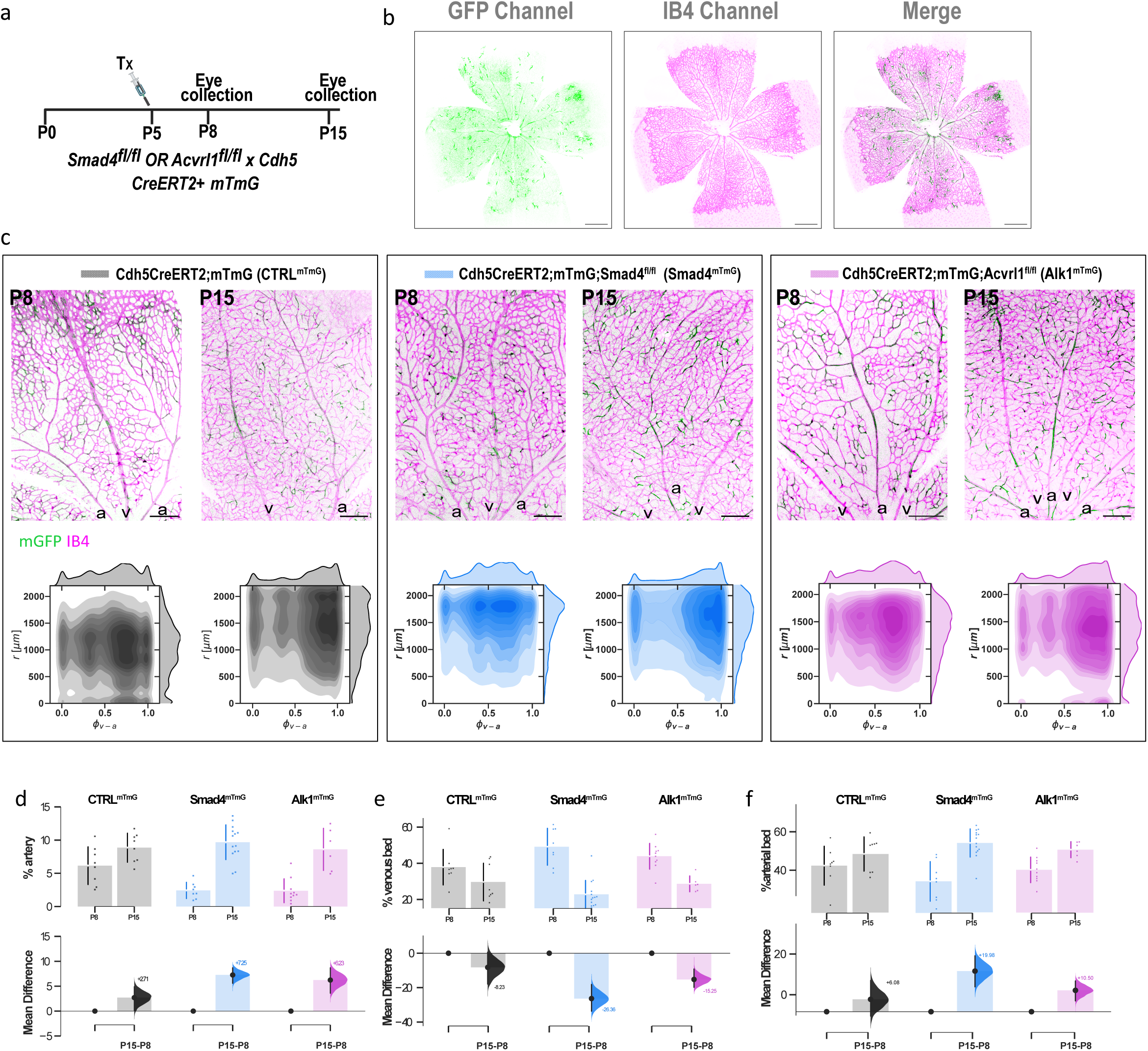
*In vivo* EC population distribution analysis. a. Tamoxifen injection regimen. Single 0.75ug injection of T-OH at P5 to induce mosaic gene deletion and subsequent expression of GFP, collection at P8 and P15. Number of retinae collected per condition: P8_CTRL^mTmG^=8; P15_CTRL^mTmG^=8; P8_Smad4^mTmG^=8; P15_Smad4^mTmG^=14; P8_Alk1^mTmG^=10; P15_Alk1^mTmG^=6; b. Representative microscopy images of P8 retina GFP, IB4 and merged channels. Scale bar - 500µm. c. Representative images of retina sections at both time points for CTRL^mTmG^ (left), Smad4^mTmG^ (middle) and Alk1^mTmG^ (right) conditions; arteries – a and veins – v. Scale bar - 250µm. Resulting KDE plots for P8 and P15 for each condition on the bottom. d. Mean difference statistical analysis of P15 GFP+EC percentage within the artery relative to P8. e. Mean difference statistical analysis of P15 GFP+EC percentage within the venous bed relative to P8. f. Mean difference statistical analysis of P15 GFP+EC percentage within the arterial bed relative to P8.

### Loss of Smad4 leads to hyper-pruning, while loss of Alk1 causes hyper-sprouting during AVM formation *in vivo*

Directional migration from vein to artery has been identified as the driving mechanism for pruning of the capillary plexus during vascular remodelling^47,48^. We therefore hypothesized that the loss of Smad4 should trigger hyperpruning, hence increased capillary regression. In contrast, reduced sensitivity and propensity to migrate against flow, as seen in Alk1 deficient ECs, should rather trigger hypopruning, causing a hyperdense plexus. In order to analyze the regression frequencies in the postnatal retinal vasculature of both Smad4 knockout (Smad4^iECKO^ hereinafter) and Alk1 knockout (Alk1^iECKO^ hereinafter), we performed immunofluorescent labeling using antibodies against CD31 and collagen type IV (ColIV) to visualize ECs and the vascular basement membrane, respectively (Fig. 6a-b). While Smad4^iECKO^ retinae overall showed a similar regression frequency to its littermate control (Fig. 6c - left, 6d – blue bars), we observed a decrease in the branching points density (Fig. 6e – blue bars), indicating elevated vessel regression at earlier time points. In contrast, we observed a significant reduction in regression frequency in Alk1^iECKO^ retinae (Fig. 6c - right, 6d – pink bars), together with increase in branching points density as well as vessel density (Fig. 6e-f). In addition, Alk1^iECKO^ retinae exclusively exhibited an overall increase in sprouting frequency, which was also visible within the AVM region (Fig. 6c, cyan triangles and supplementary Fig. 5). These results further support our hypothesis that Smad4 deletion and Alk1 deletion differentially affect remodelling of the vascular plexus.

**Figure 6:**
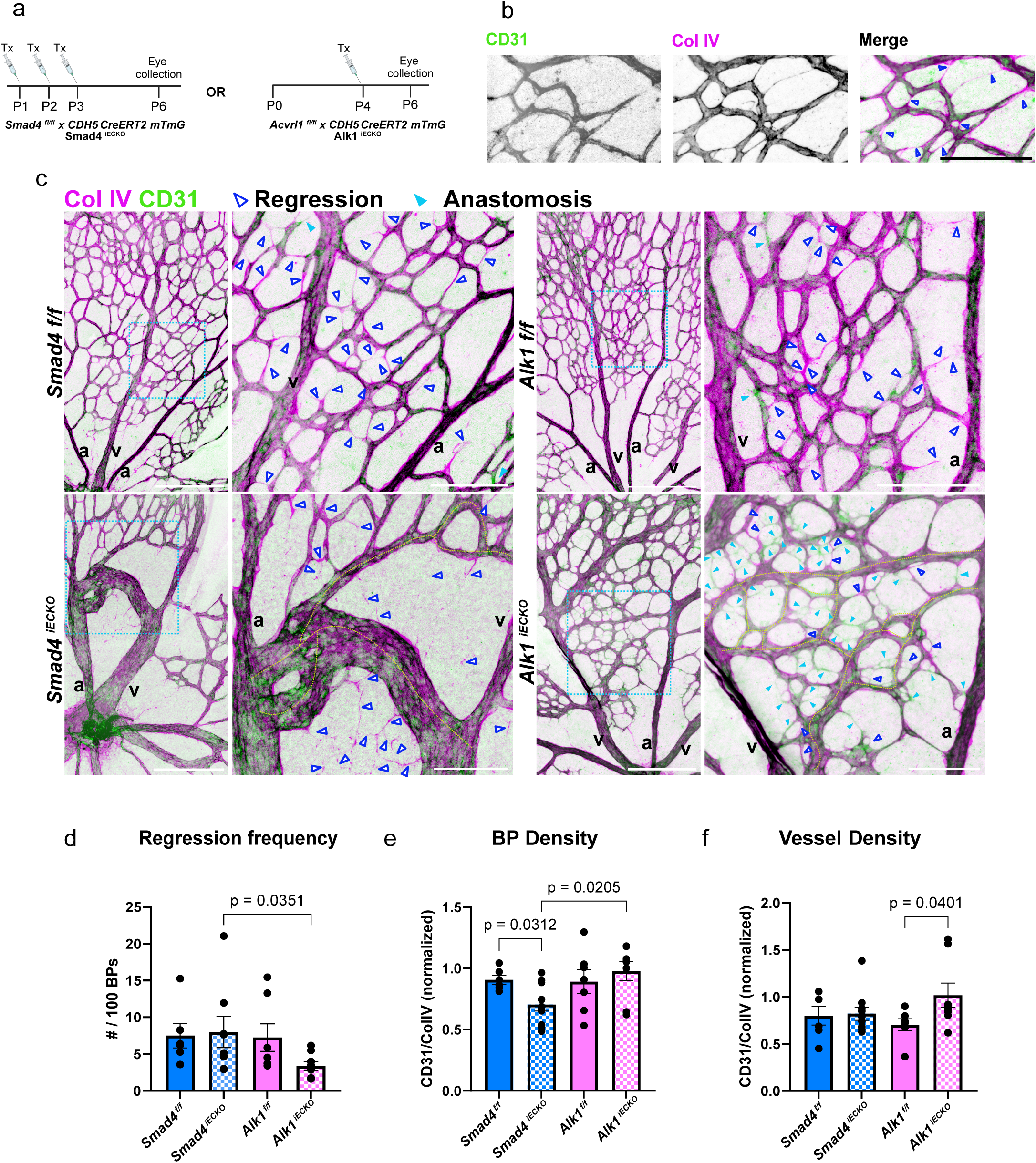
in vivo regression analysis in postnatal retina. a. Tamoxifen injection regimen for Smad4^iECKO^ and Alk1^iECKO^, respectively. N_Smad4 f/f_=6, N_Smad4 iECKO_=8, N_Alk1 f/f_=7, N_Alk1 iECKO_=8. b. Representative grayscale and merge of maximum intensity projections of CD31 channel and Col IV channel of Alk1 littermate control retina. Dark blue triangles indicate a regression event, which is defined by partial or total absence of CD31 signal within a vessel while ColIV signal is present. Scale bar 100µm. c. Representative images of immunofluorescent staining of CD31 (green) and Collagen Type 4 (magenta) in P6 littermate controls (top) and mutated (bottom) retinae. Blue rectangles annotate zoomed in sections on the right. Dark blue unfilled triangles mark regression events, filled triangles mark anastomosis events. AVMs are highlighted in yellow dashed lines. Size marker 250μm in main images, 100μm in zoomed in images; v - vein, a - artery. d. Regression frequency, presented by number of regression events per 100 CD31+ branching points. Number of branching points is extracted from the 3DVascNet software for both CD31 and Col IV channels. Regression events are counted within a region of interest (ROI) which is also used to determine the number of branching points. Two-sided non-parametric Mann Whitney test was used to determine statistical significance. Data are represented as mean ± SEM. e. Branching points’ density (# branching points/ROI volume) and f. vessel density (vessel volume/ROI volume x100%) ratios in littermate controls (full-color bars) and KO retinae (dotted bars). Two-sided non-parametrical Mann Whitney test was used to determine statistical significance. Data are represented as mean ± SEM.

## Discussion

This study identifies distinct molecular and cellular mechanisms underlying AVM formation in ECs with SMAD4 and ALK1 loss-of-function. Our findings highlight that while both SMAD4 and ALK1 loss-of-function contribute to AVM pathogenesis, the underlying cellular behaviors and transcriptomic responses differ markedly, pointing to divergent mechanisms driving these vascular pathologies.

The results reveal that SMAD4 loss-of-function enhances EC sensitivity to LSS, leading to increased hypertrophy and directional migration. The observed increase in KLF4 expression upon SMAD4 deficiency confirms recent findings^24^, lending strength to the concept that AVM formation is primarily driven by changes in EC mechanotransduction^49^. The combined effects of increasing cell size, a phenotype first reported in ENG mutants in zebrafish^50^, and our unique observation of increased directional migration, orientation and polarization against flow, suggest an intriguing pathomechanism: this heightened migratory response accelerates vascular remodelling, resulting in premature capillary regression and the formation of large, high-flow AV shunts. Such a mechanism would explain the distinct appearance of AVMs in SMAD4 mutant retinae. Our data provide several levels of evidence for increased flow sensitivity in SMAD4 deficient ECs: First, transcriptional changes highlight besides classical flow-responsive genes like KLF4 and KLF2, also prominent flow-regulated upregulation of cell surface proteoglycans involved in endothelial permeability regulation and vessel lumen regulation, like PODXL^51–54^ (Supplementary Fig. 1.3a). Second, flow-mediated cell shape changes and directional polarity are surprisingly enhanced rather than diminished in the absence of SMAD4 (Figs 2 and 3). Third, directional EC population movements both *in vitro* (Fig. 4) and *in vivo* identify increased movement towards the artery, a behaviour depending on flow-responses^9^ (Fig. 5). The cause for such enhanced flow sensitivity in SMAD4-deficient ECs remain to be identified.

One possibility is that ALK1 and SMAD4 differentially tune endothelial sensitivity to pro-angiogenic versus remodelling cues. Recent work by Franco and colleagues’ identified competition between the chemotactic gradient of VEGFA and the mechanical directional response due to flow^7^. Conceptually, this competition plays out in the capillary transition zone where VEGFA gradients and flow gradients compete. Intriguingly, both processes involve VEGFR signaling, with evidence for ligand-independent activation of VEGFR2 by flow, versus ligand-dependent activation in the context of sprouting. Furthermore, Notch signaling is linked to controlling arterial and venous cell fate acquisition^55,56^, sprouting^57^ as well as flow-induced remodelling^58–60^. Thus, the loss of ALK1 may drive hypersprouting and venous/capillary overgrowth by reduced Notch and increased ligand-dependent VEGFR2 signalling, whereas the loss of SMAD4 selectively enhances Notch related pruning through flow-dependent VEGFR2 mechanisms. In line with this idea, GO term analysis highlights a consistent pattern of transcriptional dysregulation in VEGFR signaling and Integrin mediated cell adhesion in *siSMAD4*-treated ECs relative to *siALK1*-treated ECs (Fig. 1c, d and supplementary Fig. 1.2, 1.3): Gene sets involved in negative regulation of VEGFR signaling (M22989, M25085) show predominant upregulation of their constituent genes, while positive regulators (M15928) exhibit downregulation. In addition, GO terms related to integrin-mediated cell adhesion showed a predominance of upregulated genes (M23222, M23221, M13232). These data suggest a transcriptional reprogramming that may reduce VEGFR2 pathway activity, potentially altering cellular responses to flow stimuli. As VEGFR2 activation can occur via ligand-dependent (VEGF-A-mediated) or ligand-independent (flow-induced) mechanisms^61^, shifts in the expression of key regulatory genes may influence the dominant signaling mode. In *siSMAD4*-treated cells, DPP4 and PODXL were enriched as part of integrin-mediated cell adhesion GO Terms, and were uniquely upregulated - DPP4 in a flow-independent manner, and PODXL in a flow-dependent manner, relative to both *siCTRL*- and *siALK1*-treated cells (Fig. 1d, Supplementary Fig. 1.3a, b). Both genes are implicated in modulating endothelial adhesion and migration^53,62,63^, and may contribute to the altered flow responsiveness observed. DPP4 in particular, has been linked to endothelial-to-mesenchymal transition (EndMT) through a TGF-β2-dependent, SMAD-independent mechanism^64^. Its interaction with integrin β1 has been shown to upregulate VEGFR1 expression, which acts as a decoy receptor and suppresses VEGFR2 signaling. In parallel, we observed downregulation of ITGAV and ITGB3 – integrins that facilitate VEGF-A binding and ligand-dependent VEGFR2 activation (Supplementary Fig.1.3c,d)^65^. This combination suggests a shift away from VEGF-A-dependent signaling and toward ligand-independent, flow-induced VEGFR2 activation, which may underlie the enhanced sensitivity to flow observed in SMAD4-deficient ECs. However, further experimental validation is required to confirm these regulatory mechanisms and their functional consequences.

This molecular shift is mirrored in the striking vascular phenotype observed *in vivo*. SMAD4 loss-of-function in the postnatal mouse retina leads to large shunts devoid of a surrounding capillary plexus. Yet, even the peripheral plexus that will be hypoxic, showing upregulation of hypoxia induced genes, like ANGPT2^66^, shows no sign of hypersprouting, unlike in the case of ALK1 mutant conditions. Thus, the lack of SMAD4 appears to selectively increase endothelial responses to flow, whilst reducing endothelial responses that are normally triggered by hypoxia. This phenotypic divergence reinforces the notion that SMAD4 plays a key role in balancing competing angiogenic cues and may act as a molecular switch between sprouting- and flow-mediated endothelial programs.

In contrast, ALK1 loss-of-function appears to impair endothelial sensitivity to LSS, disrupting vascular remodelling and leading to reduced capillary regression (Fig. 6). We observed a blunted upregulation of KLF4 and diminished responses to LSS across multiple parameters, including reduced polarization, orientation (Fig. 2,3) and migration of ALK1-deficient cells both *in vitro* and *in vivo* (Fig. 4,5). This remodelling failure results in hypervascularized networks with dense, multi-capillary AVMs (Fig. 6d–f). While directional migration is impaired, it does not appear to be the primary mechanism driving AVM formation: in mosaic monolayers under flow, *siALK1* cells show reduced directional migration during the first half of the experiment. However, in the second half, both *siCTRL* and *siALK1* populations migrate against the flow direction - doing so even earlier than cells in *siCTRL-*mosaic cultures. Similarly, in mosaic retinae, ALK1-deficient GFP⁺ ECs still migrate toward the artery, albeit less efficiently. These findings indicate that although ALK1 deficiency delays flow-induced migration, it does not completely abolish the endothelial capacity to migrate in response to mechanical cues.

Instead, our transcriptional data point to a more fundamental defect in endothelial fate specification and quiescence that arises prior to flow onset. ALK1-deficient ECs already show downregulation of key fate regulators, including the arterial markers GJA4 (Connexin37) and CDKN1B (p27) (Supplementary Fig. 1.3e,f)^55^, as well as the venous markers BMP4^8^ and NR2F2 (COUP-TFII)( (Supplementary Fig. 1.3g,h)^56,67^. In addition, Notch signaling appears uncoupled from flow stimuli: canonical Notch targets DLL4, HEY1, and HEY2 do not respond appropriately under shear stress, particularly at 4h (Supplementary Fig. 1.3i-k). These early changes coincide with persistent ANGPT2 upregulation (Supplementary Fig. 1.3l) and downregulation of its receptor TEK (Tie2) (Supplementary Fig.1.3m), even before the onset of flow. Given that ANGPT2 is a potent pro-angiogenic factor normally suppressed under flow to promote quiescence^68,69^, this pattern suggests that ALK1-deficient ECs remain in an immature, angiogenic-like state – unable to transition into a flow-responsive phenotype. Our findings are supported by recent work in ALK1-null hiPSC-derived ECs, where ANGPT2 upregulation and reduced pSMAD1/5/9 signaling similarly marked a persistent angiogenic state^70^.

While previous studies comparing phenotypes and pathomechanisms of AVM formation between SMAD4, ALK1 and ENG mutants highlighted similarities, including upregulation of ANGPT2^71^, we only find ANGPT2 upregulation in ALK1 deficient cells, in line with the notion of failed induction of quiescence^68,72^. A possible reason for this apparent discrepancy is the fact that we analyzed cell-autonomous gene regulation in ECs under flow, in the absence of pathology. Once an AVM has formed *in vivo*, studying altered gene regulation will report changes that may be primary, but also changes that are secondary to the altered tissue environment, including hypoxia. ANGPT2 is a canonical marker of endothelial activation by hypoxia, but also in inflammatory conditions^69,73^. The fact that large areas of the endothelium upregulate ANGPT2 in bulk RNAseq of isolated ECs from Smad4 mutant retinae^66^, like those from Alk1 mutant retinae, is not necessarily evidence for a common causative ANGPT2-induced pathomechanism. Nevertheless, ANGPT2 overexpression may contribute as a driver of wider tissue responses in both conditions.

Our findings challenge the notion that AVM formation in SMAD4 and ALK1 loss-of-function follows a unified pathway. Instead, they support a model in which SMAD4 loss-of-function promotes FSS-driven vascular remodelling, while ALK1 loss-of-function primarily disrupts cell fate acquisition and exit from an angiogenic state. These insights align with prior studies implicating BMP signaling dysregulation in AVM pathogenesis^26,57,74^, but also highlight the necessity of pathway-specific therapeutic strategies.

This study opens several avenues for future research. The observed differences in transcriptional and behavioral responses between SMAD4 and ALK1 loss-of-function highlight the need for further mechanistic studies using multi-omics approaches. Combining transcriptomics, proteomics, and single-cell analyses across various tissue contexts could clarify which signaling effectors are primary drivers of AVM formation versus secondary responses to established pathology. Additionally, exploring factors such as ANGPT2 in ALK1 loss-of-function and DPP4 in SMAD4 loss-of-function could uncover novel targets for modulating EC behavior.

In conclusion, this work provides a mechanistic framework for understanding AVM pathogenesis and underscores the need for pathway-specific therapeutic strategies in addressing vascular anomalies such as HHT.

## Supporting information

supplemental figures merged

## Non-standard Abbreviations and Acronyms

AVM: Arteriovenous Malformation
HHT: Hereditary Hemorrhagic Telangiectasia
BMP: Bone Morphogenetic Protein
EC: Endothelial Cell
FSS/LSS/HSS: Fluid/Low/High Shear Stress
siRNA: Small Interfering RNA
siCTRL: Control siRNA
siSMAD4: siRNA against SMAD4
siALK1: siRNA against ALK1
GO_BP: Gene Ontology – Biological Process
DEG: Differentially Expressed Gene
VEGF/VEGFR: Vascular Endothelial Growth Factor/ Vascular Endothelial Growth Factor Receptor
DPP4: Dipeptidyl Peptidase-4
PODXL: Podocalyxin
TEK: TEK Tyrosine Kinase (encodes Tie2)
ENG: Endoglin
ALK1: Activin Receptor-Like Kinase 1 (ACVRL1)
SMAD4: Mothers Against Decapentaplegic Homolog 4
KLF4: Krüppel-like Factor 4
pSMAD159: Phosphorylated SMAD1/5/9
GJA4: Gap Junction Alpha-4 Protein (encodes Connexin 37)
CDKN1B: Cyclin-Dependent Kinase Inhibitor 1B (encodes p27)
NR2F2: Nuclear Receptor Subfamily 2, Group F, Member 2 (encodes COUP-TFII)
DLL4: Delta-Like Ligand 4
HEY1/HEY2: Hairy/Enhancer-of-split related with YRPW motif protein 1/2
ANGPT2: Angiopoietin-2
ColIV: Collagen Type IV
CD31: Cluster of Differentiation 31 (PECAM-1)
ROI: Region of Interest
DABEST: Data Analysis with Bootstrap-Coupled Estimation
HUVEC: Human Umbilical Vein Endothelial Cell
mTmG: Membrane Tomato–Membrane GFP reporter
T-OH: Tamoxifen
PI: Polarity Index
V-score: Signed Polarity Index (for directionality)
NUC/Cyt: Nucleus-to-Cytosol (ratio)
P: Postnatal Day (e.g., P4, P5, P8)

## Acknowledgements

We thank Leducq ATTRACT network members Hemaxi Narotamo and Claudio Franco, Douglas Marchuk, David Hardman and Miguel Bernabeu for providing ongoing support throughout this project, as well as Tatiana Borodina, Eireen Bartels-Klein, Irene Hollfinger for technical assistance and guidance.

## Author contributions

Olya Oppenheim: Conceptualization; data curation; formal analysis; investigation; methodology; software; project administration; validation; visualization; writing – original draft; writing – review and editing.

Wolfgang Giese: Formal in vitro analysis; methodology; software; visualization; writing – review and editing.

Hyojin Park: In vivo data curation; investigation; methodology; validation; writing – review.

Elisabeth Baumann: Formal in vivo analysis; methodology; software; visualization.

Andranik Ivanov: Computational analysis of the transcriptomic data; methodology; writing – review and editing.

Dieter Beule: Project administration; funding acquisition.

Anne Eichmann: Conceptualization; funding acquisition; investigation; writing – review.

Holger Gerhardt: Conceptualization; investigation; funding acquisition; supervision; project administration; writing – review and editing.

## Data Accessibility

The raw RNA-Seq reads are deposited in the GEO database under accession number GSE282952.

## Supplementary figure legends

**Supplementary figure 1:**

a. Knockdown verification of SMAD4, ALK1. Top – qPCR analysis of *siCTRL* (black), *siSMAD4* (blue), *siALK1* (magenta), *siDouble* (lime), 48h post siRNA treatment. Log2FC values represent relative expression to b-actin. N=3 independent experiments. Bottom – Western blot analysis of *siCTRL*, *siSMAD4* and *siALK1* with representative images of blots, 48h post siRNA treatment. Bands normalized to GAPDH; n=3 independent experiments.
b. Full panel plot of GO_BP of *siSMAD4* vs. *siALK1* in static, 4h and 16h of LSS. GO_BPs are clustered according to their enrichment in the different flow conditions: Cluster I – enrichment in all flow conditions; Cluster II – enrichment in static only; Cluster III – enrichment in 4h LSS only; Cluster IV – enrichment in 16h LSS only; Cluster V – enrichment in both 4h and 16h LSS; Cluster VI – enrichment in static and a 4h LSS **OR** 16h LSS.
c. Expression patterns of individual genes over time (RNAseq data). Y axis is Log2FC scale, X axis are flow conditions. Expression patterns of a. PODXL, b. DPP4, c. ITGAV, d. ITGB3, e. CDKN1B (p27), f. GJA4 (Cx37), g. BMP4, h. NR2F2, i. DLL4, j. HEY1, k. HEY2, l. ANGPT2, m. TEK (Tie2). *siCTRL* – black; *siSMAD4* – blue; *siALK1* – magenta.

**Supplementary figure 2:**

Representative pSMAD159 immunofluorescence images of HUVECs after exposure to HSS (1.8Pa). Scale bar 100μm.

**Supplementary figure 3:**

a. Representative KLF4 immunofluorescence images of HUVECs after exposure to static, 4h and 16h LSS conditions. Scale bar 100μm.
b. Representative VEcad staining of HUVECs after exposure to static, 4h and 16h of LSS conditions. Scale bar 100μm.
c. –e Mean difference statistical analysis of cell area (c), cell elongation (d) and cell orientation (e) of *siCTRL-*, *siSMAD4-* and *siALK1*-treated cells at static, 4h and 16h of LSS. The further away the difference histogram is from the control group line (0.0), the higher the significance.

**Supplementary figure 4:**

Mosaic EC population shift *in vivo*.

a. Analysis overview: two-dimensional masks of retina outline, arteries and veins in flat-mounted retinas, result in a 2D map. Each GFP+ EC’ coordinates are based on its location relative to the vein(0.0) and artery (1.0) and the radial distance from the optic nerve to the sprouting front. Kernel Density Estimate (KDE) plots represent the cumulative location of all GFP+ EC across all retinae for each condition.
b. Density histograms of CTRL^mTmG^ (black), Smad4^mTmG^ (blue) and Alk1^mTmG^ (magenta) in overlay, describing GFP+EC distance relative to artery at P8 (left) and P15 (right).
c. Mean difference statistical analysis of P8 (left) and P15 (right) GFP+EC distribution in the 75^th^ percentile on the vein-artery axis, equivalent to the peri-arterial capillary bed.
d. Mean difference statistical analysis of P8 (left) and P15 (right) GFP+ EC distribution the 90^th^ percentile on the vein-artery axis, equivalent to the arteries and immediate surrounding vessels.

**Supplementary figure 5:**

Grayscale Col IV and CD31 channel images of enlarged sections of Figure 6. Scale bar 100 µm.

## Supplementary movies

Live imaging of endothelial migration under flow.

**1.** Representative time lapse imaging of HUVECs labeled with SPY505 and exposed to LSS for 48h.

a. siCTRL; b. siSMAD4; c. siALK1. Frame interval – 7.5 minutes.

**2.** Representative time lapse imaging of HUVECs labeled with CellTracker Green (siCTRL) and CellTracker Red (siCTRL/siSMAD4/siALK1) and exposed to LSS for 62h. a. siCTRL mosaic (green + red); b. siCTRL green+siSMAD4 red; c. siCTRL green+siALK1 red.

