## supplemental figures merged for "Gene-specific endothelial programs drive AVM pathogenesis in SMAD4 and ALK1 loss-of-function"

Supplementary figure 1.1

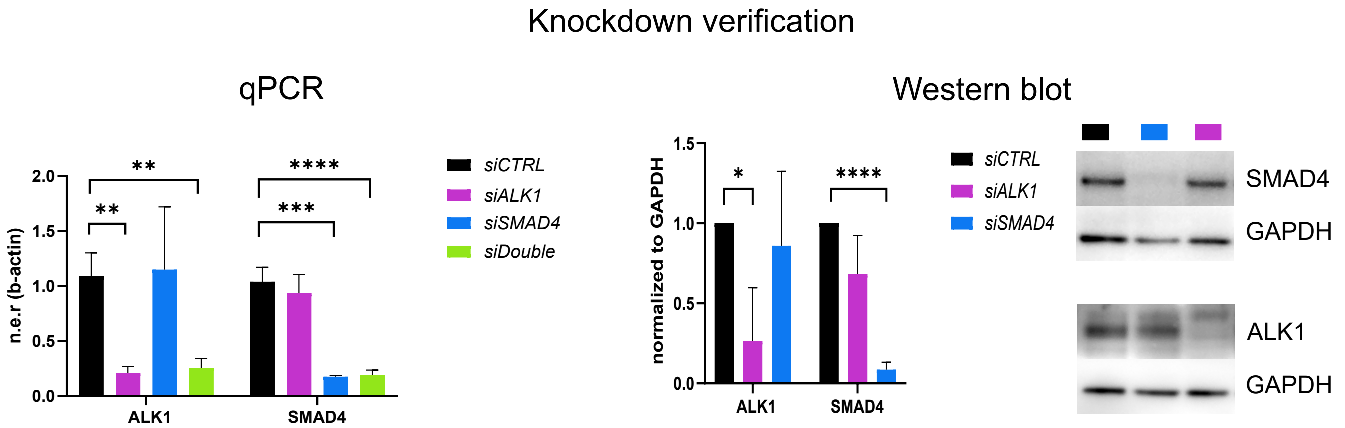

Supplementary figure 1.2

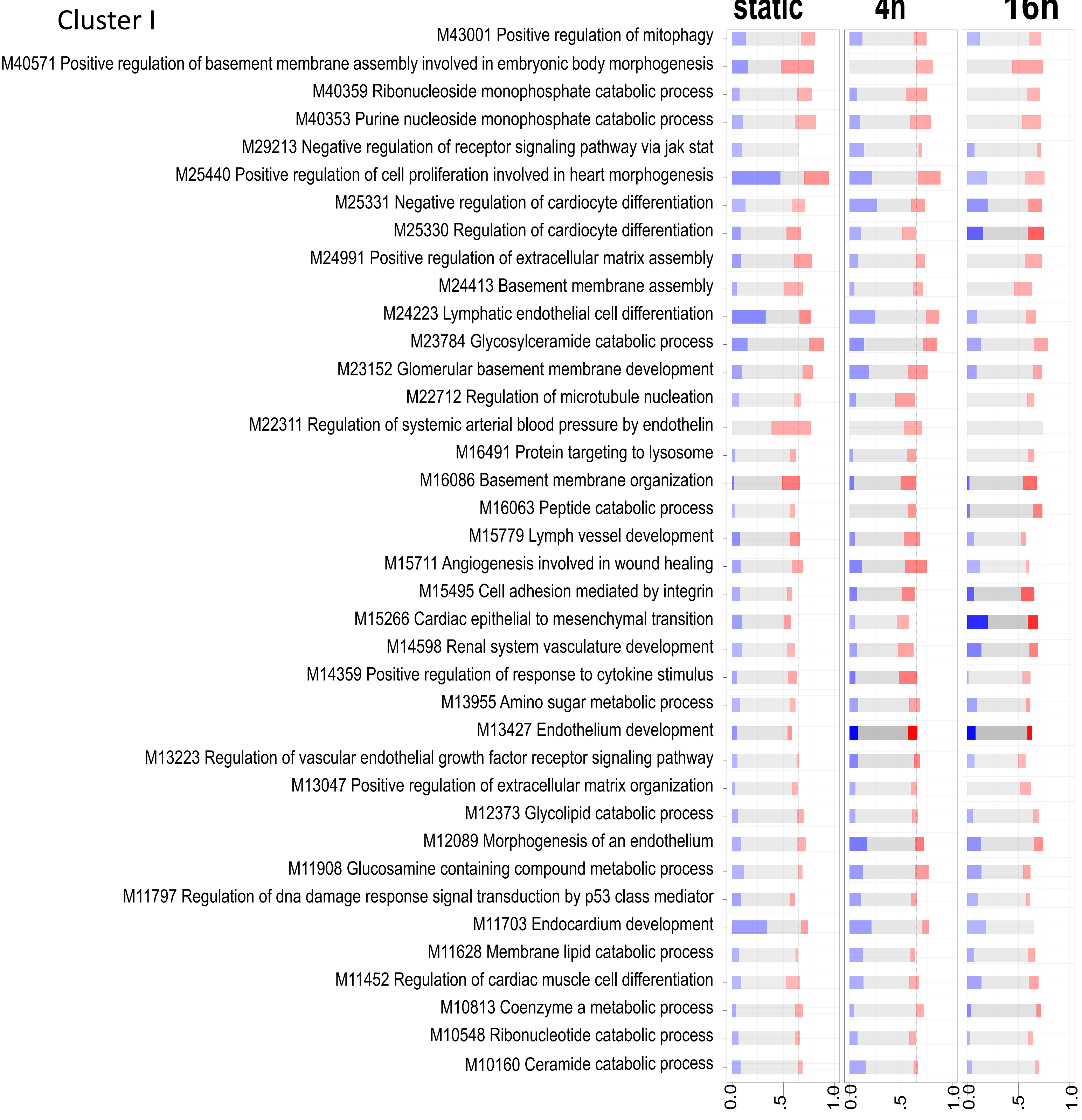


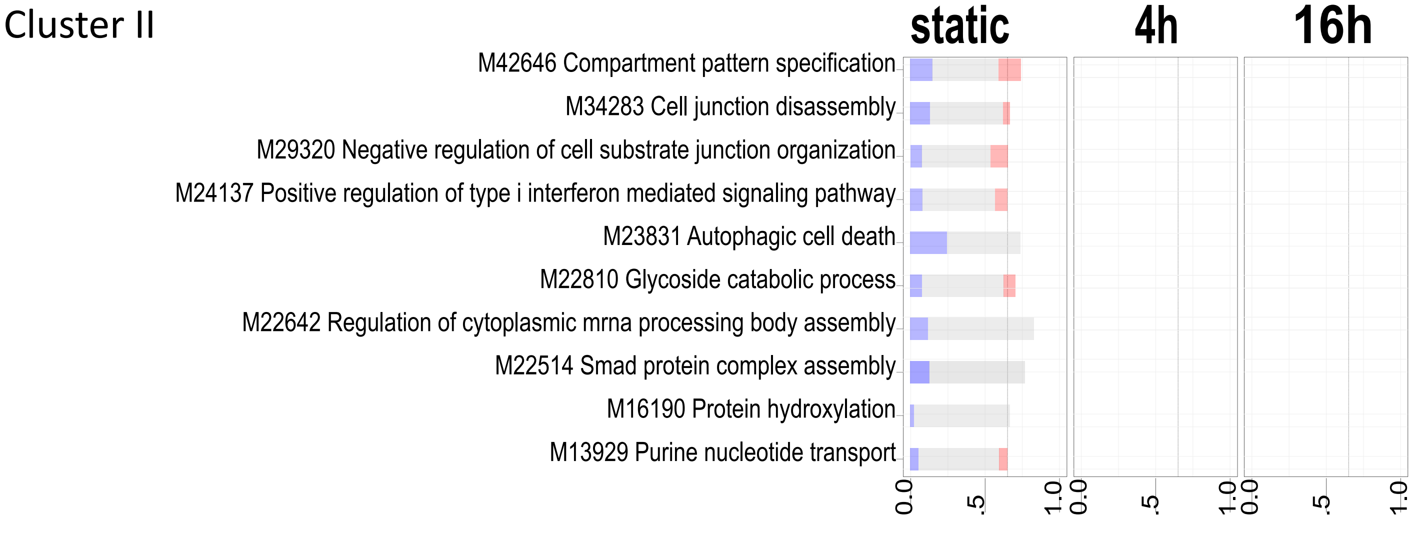

Supplementary figure 1.2 - continued

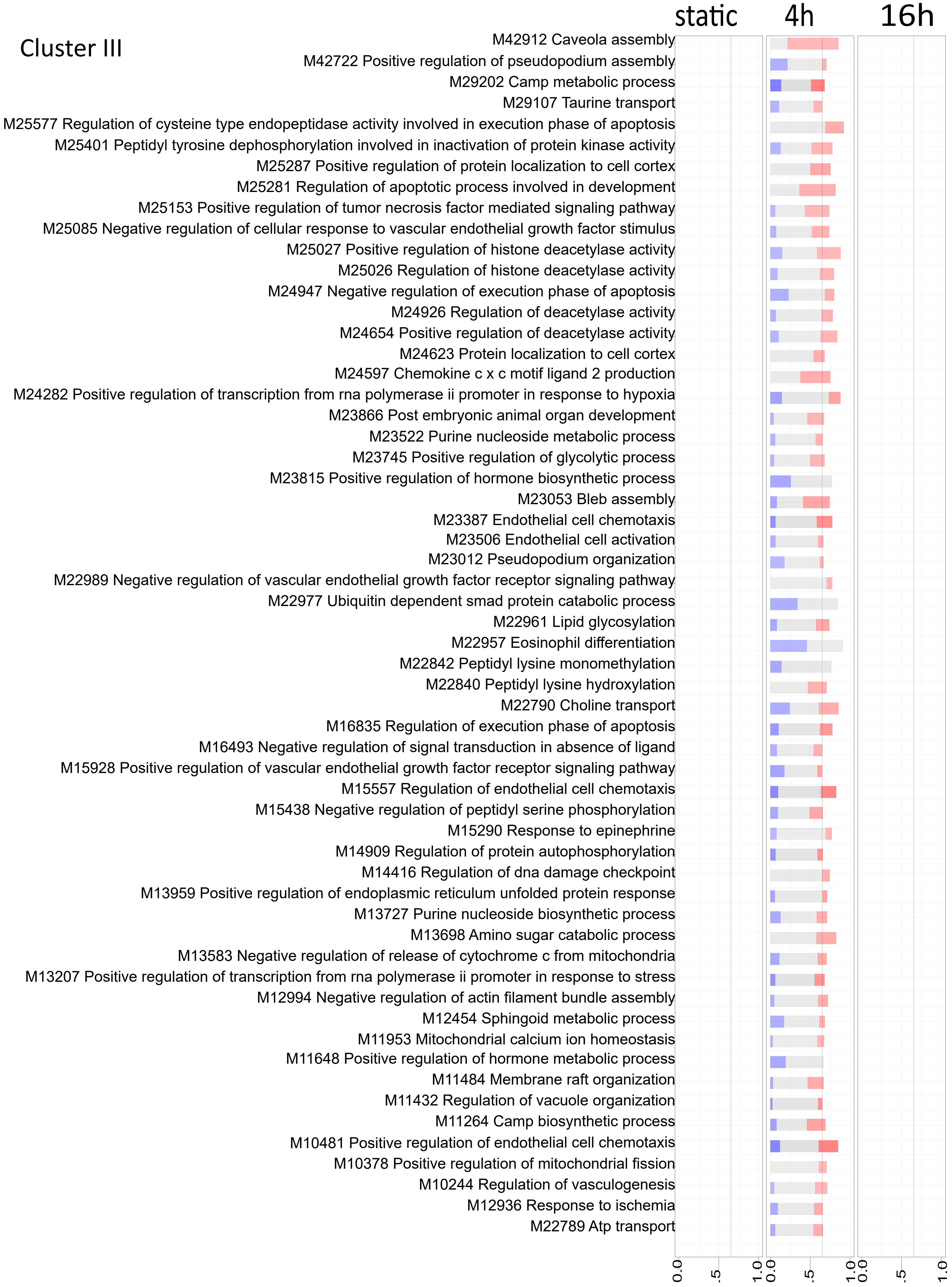

Supplementary figure 1.2 - continued

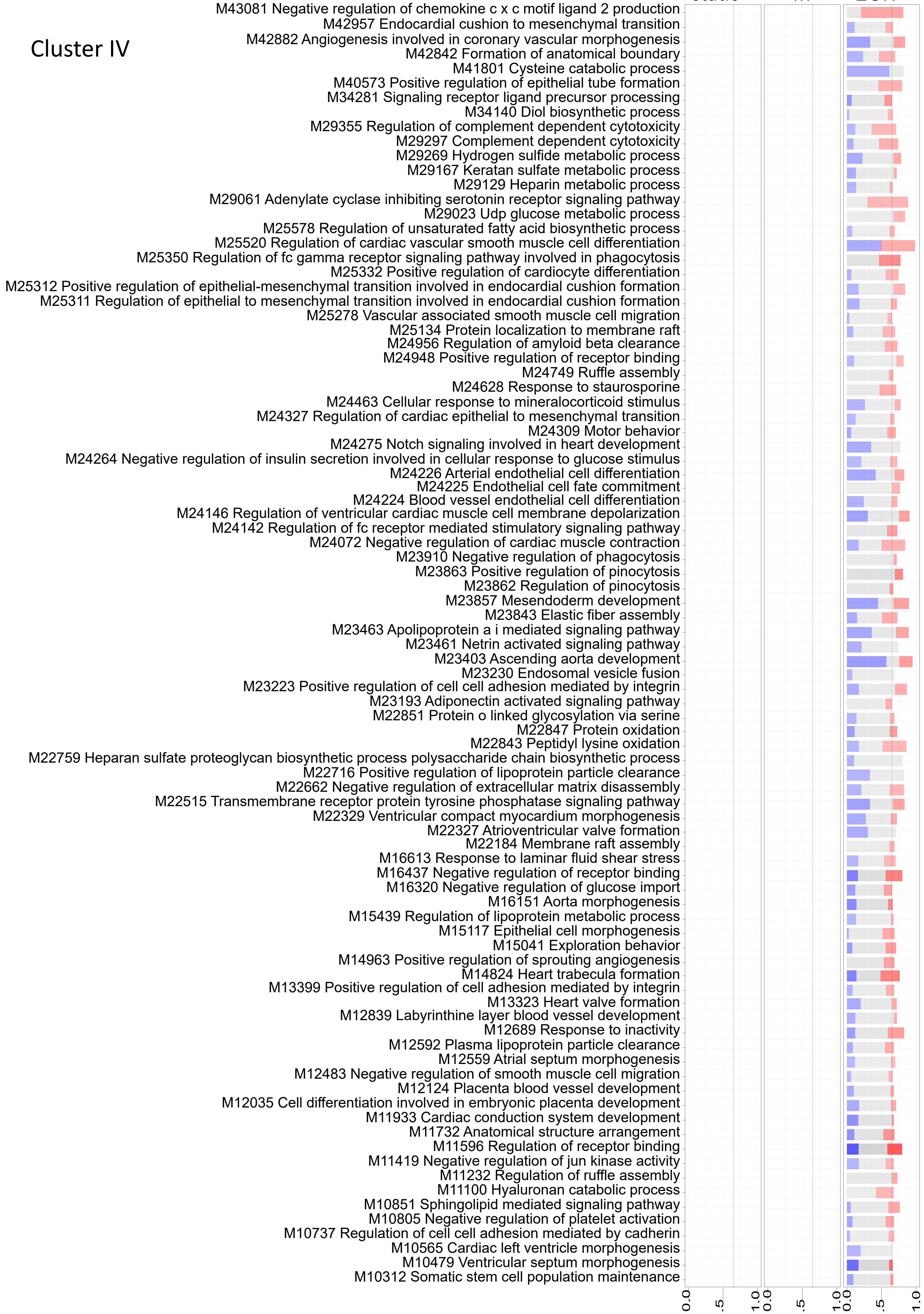

Supplementary figure 1.2 - continued

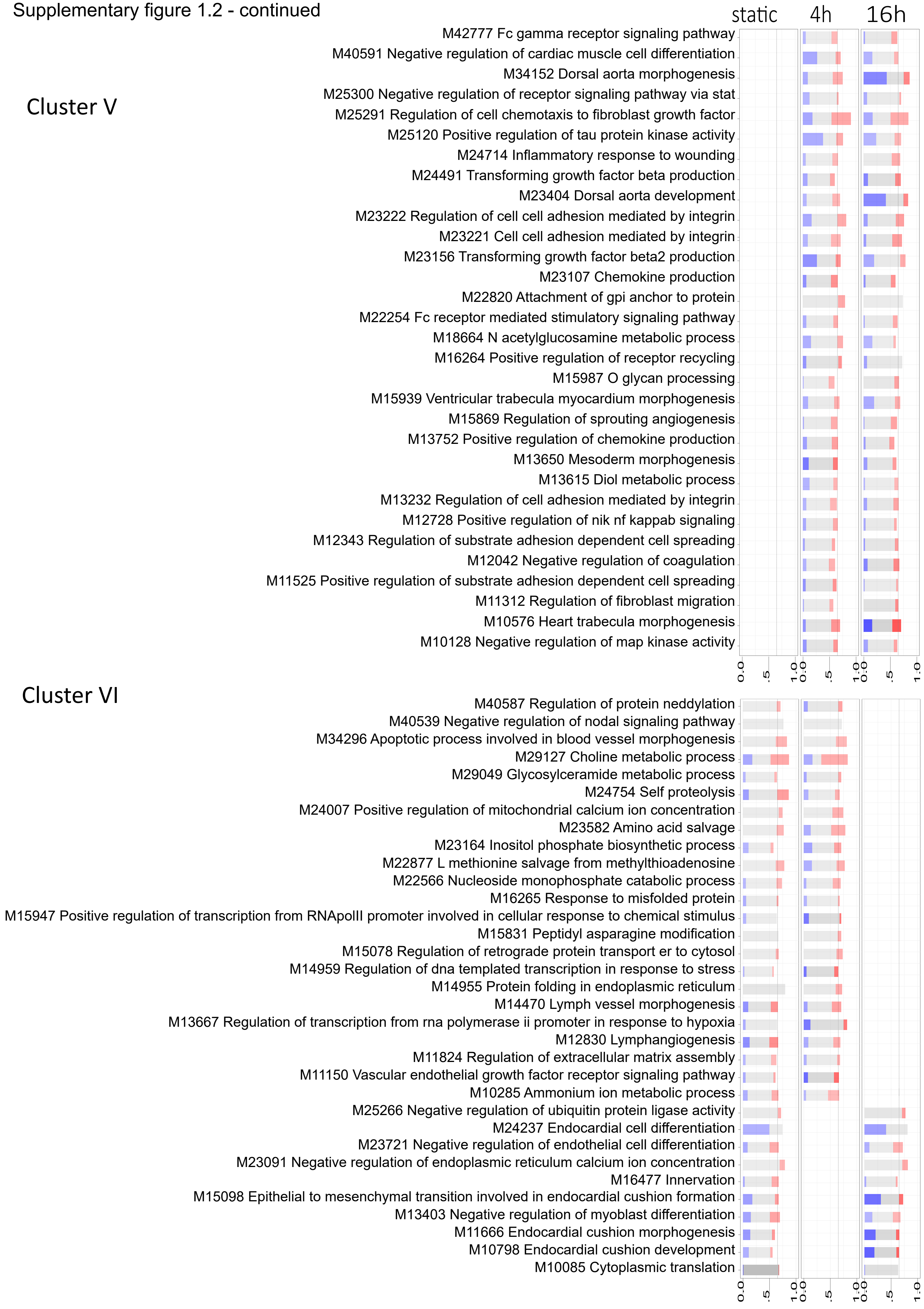

Supplementary figure 1.3

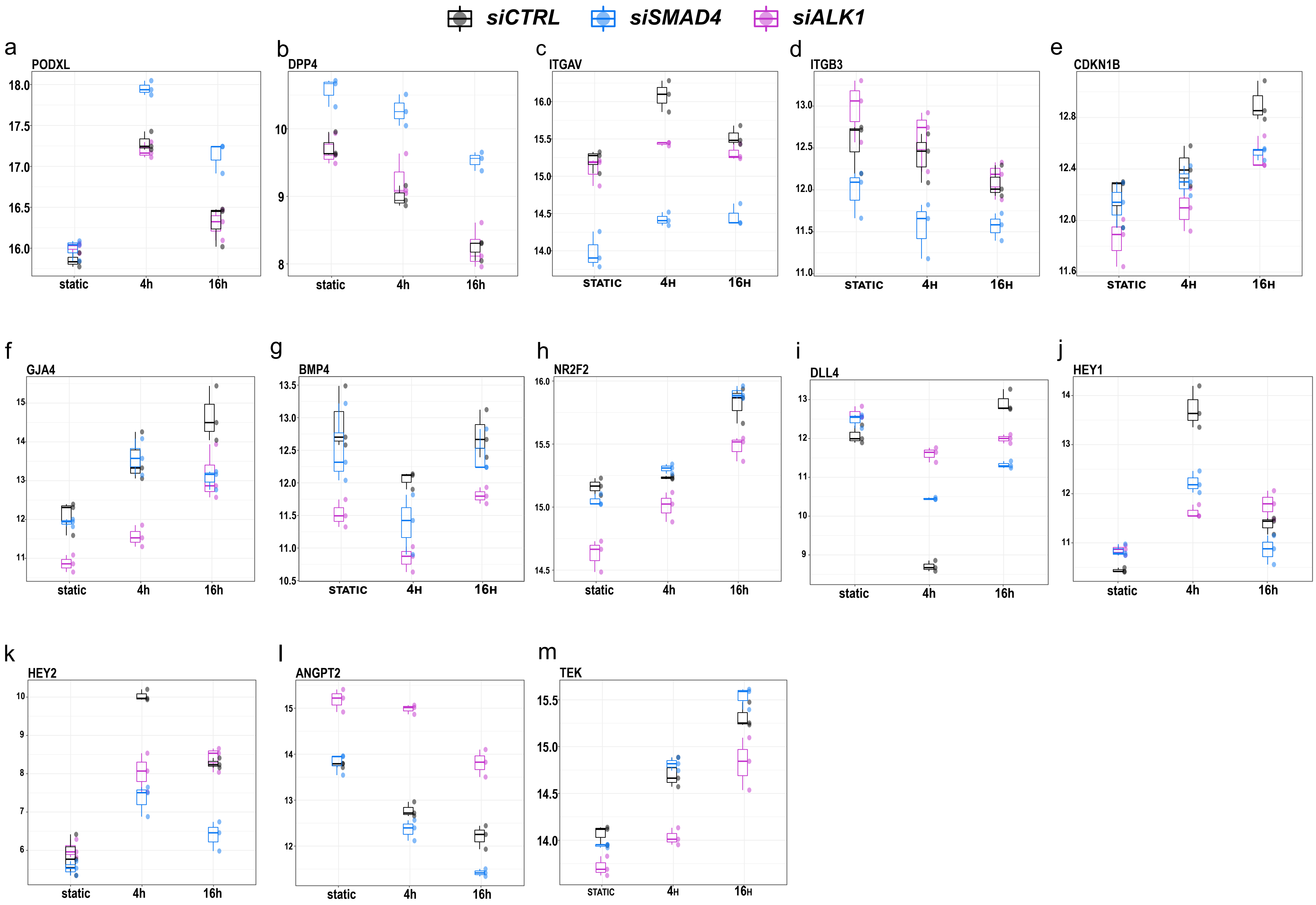

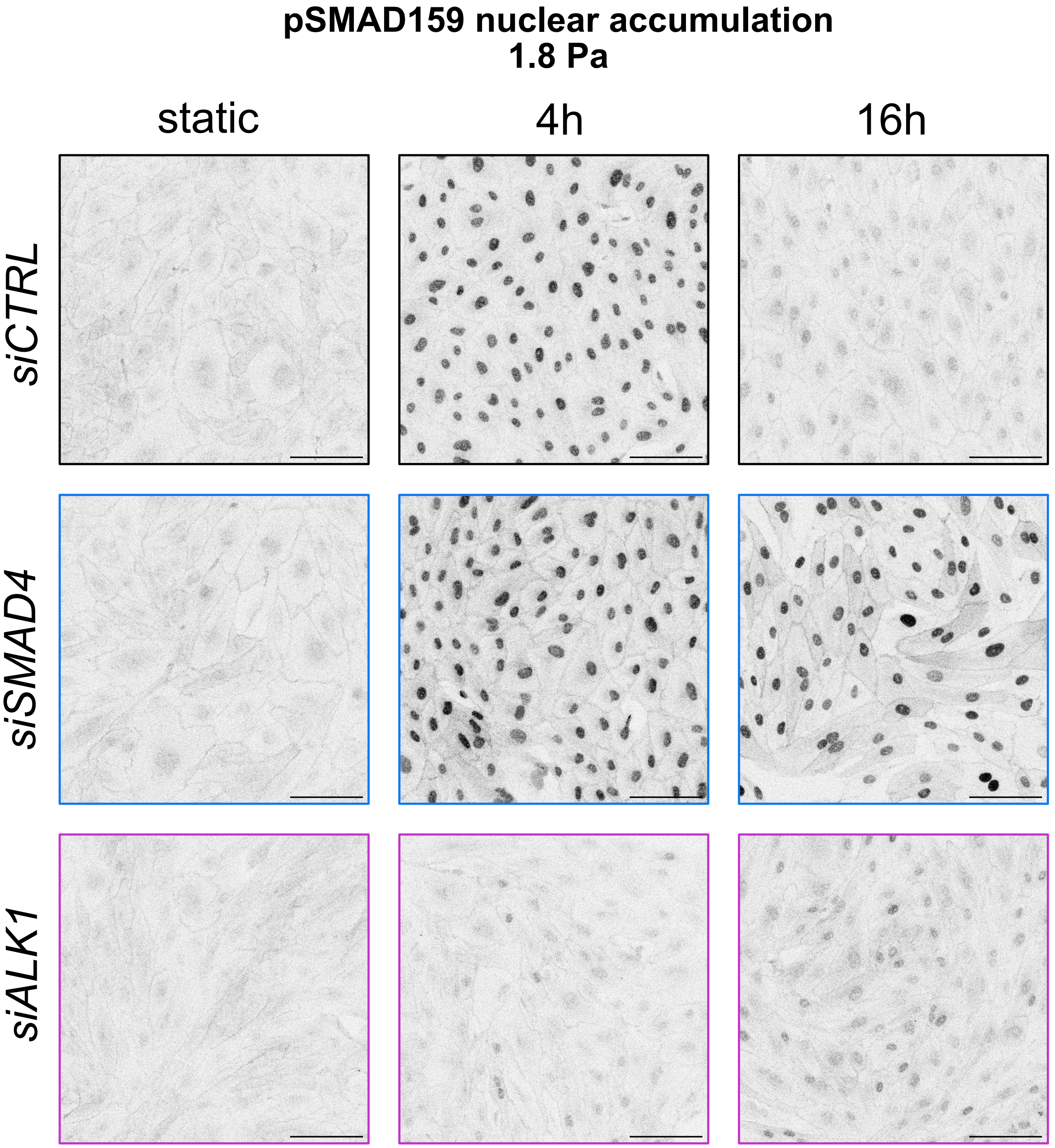
pSMAD159 nuclear accumulation
1.8 Pa
static
4h
16h
siCTRL
siSMAD4
siALK1

Supplementary figure 3

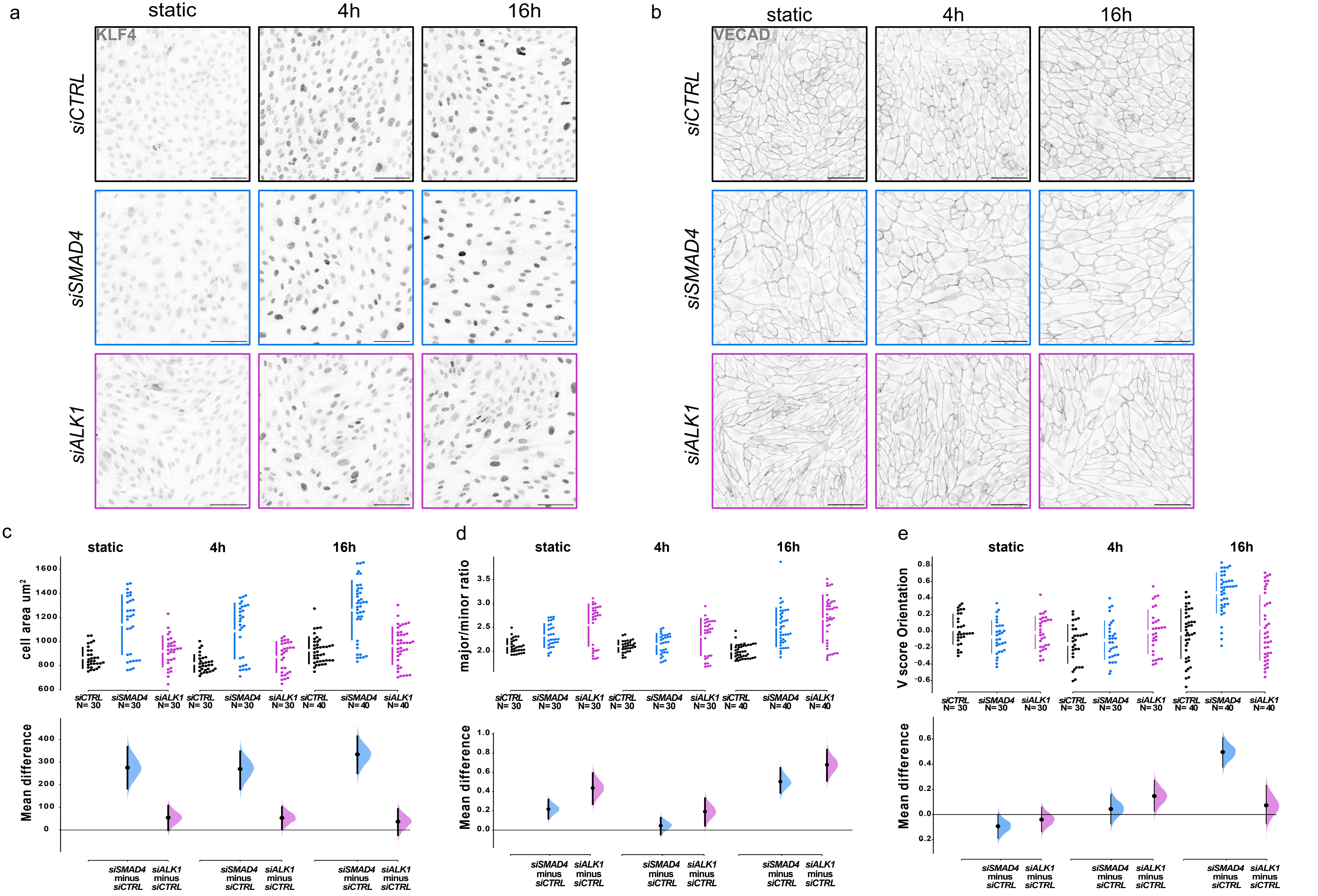

Supplementary figure 4

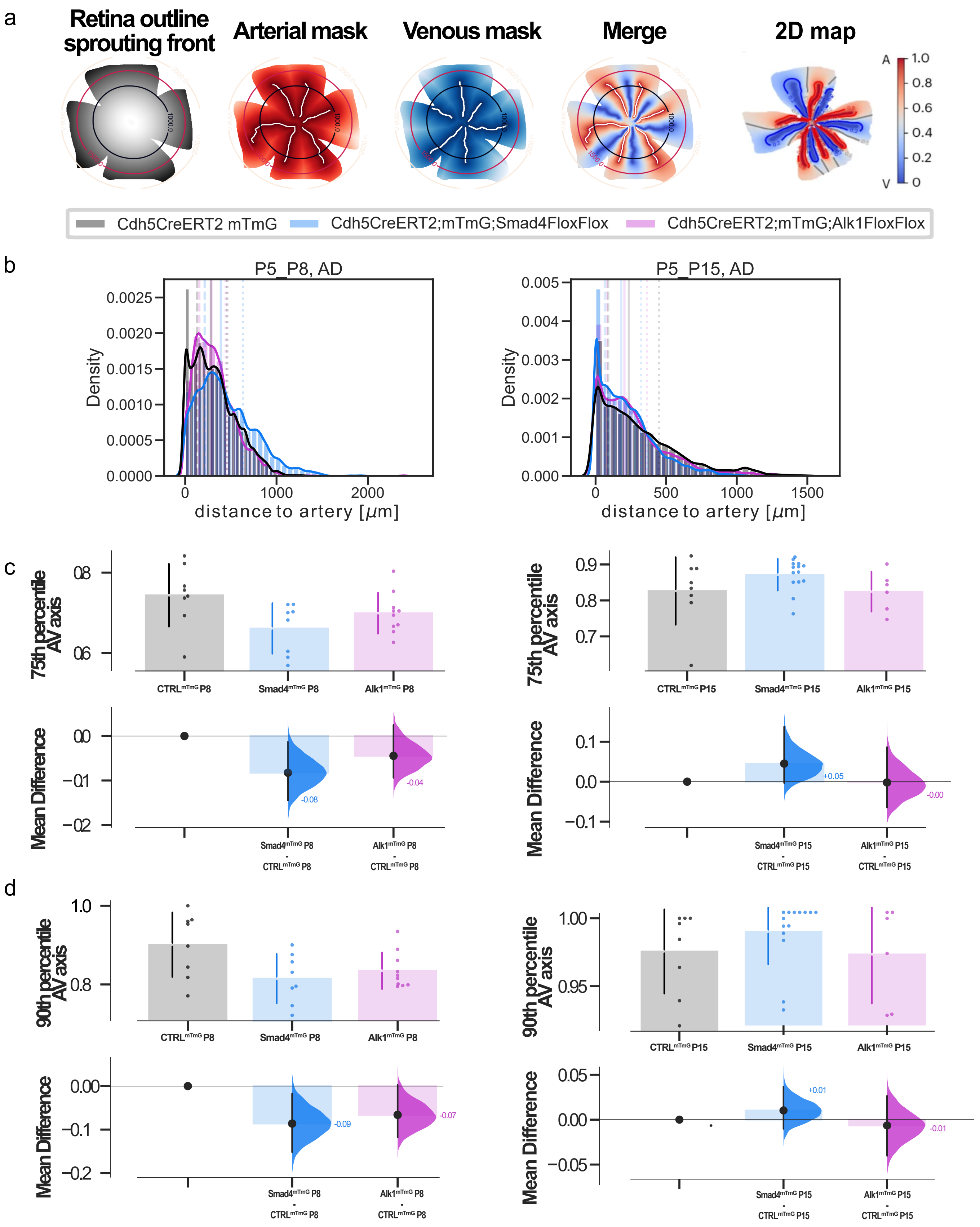

Supplementary figure 5

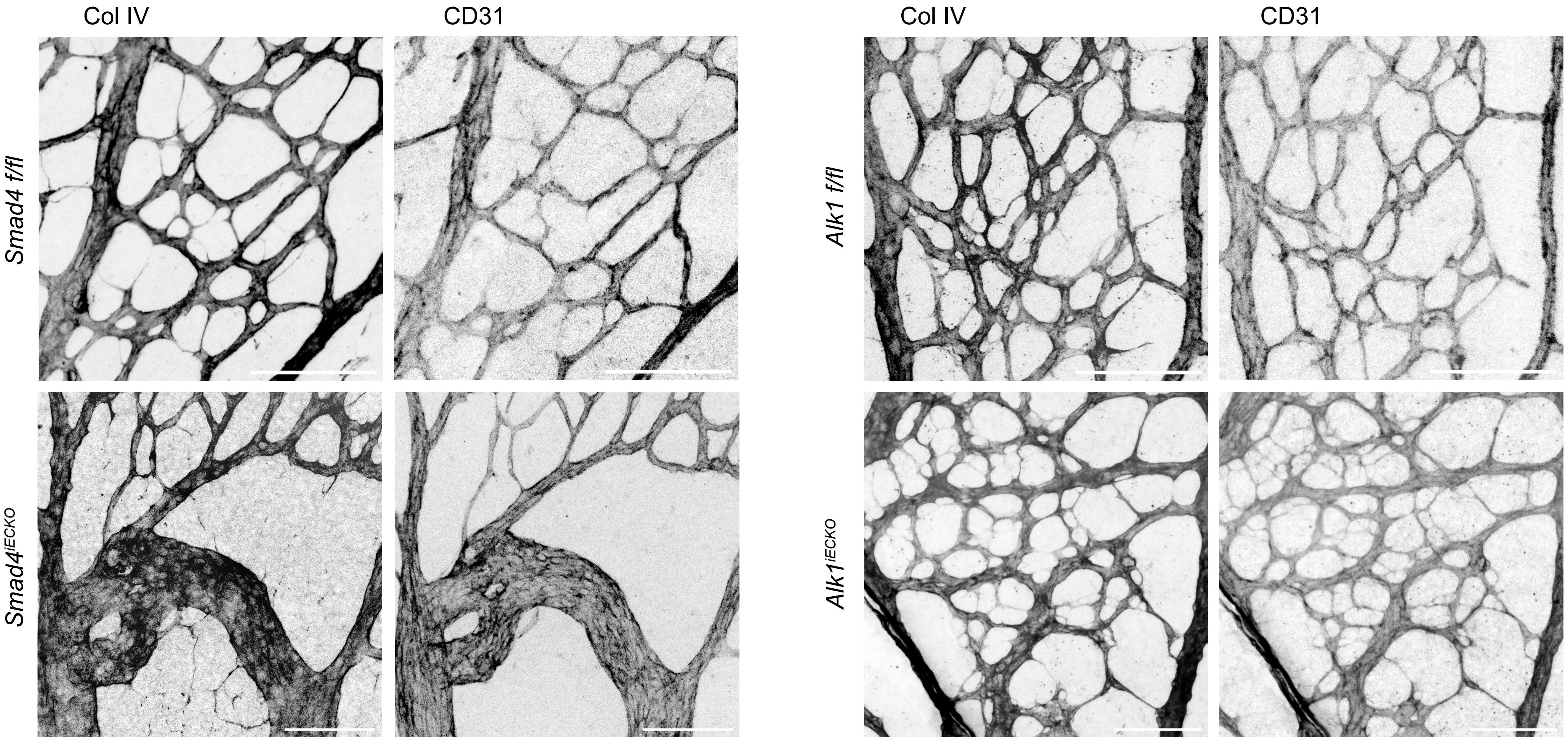
