## supplementary movie 1 for "Gene-specific endothelial programs drive AVM pathogenesis in SMAD4 and ALK1 loss-of-function"

### Slide 1
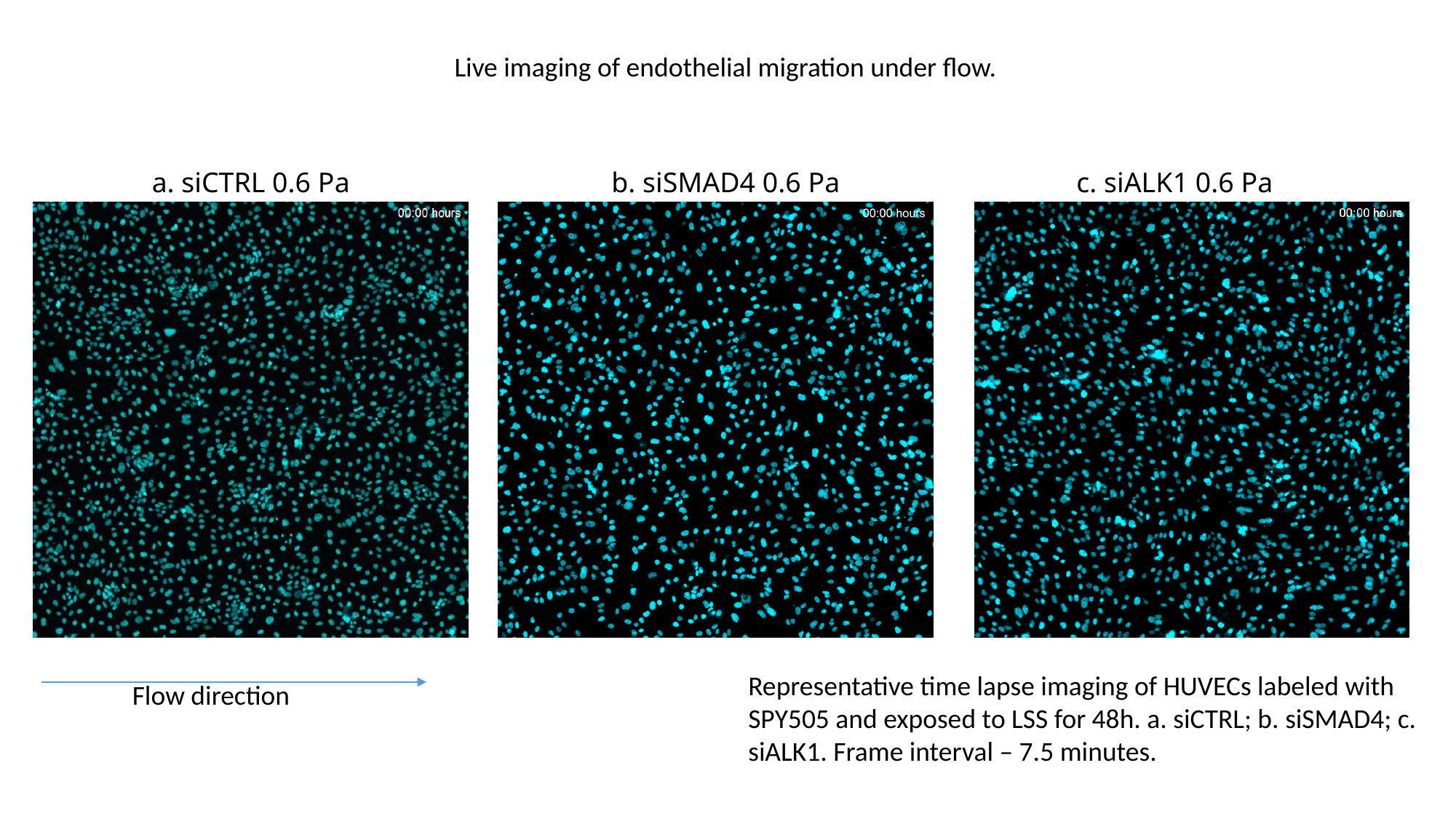

Live imaging of endothelial migration under flow.
a. siCTRL 0.6 Pa
b. siSMAD4 0.6 Pa
c. siALK1 0.6 Pa
Representative time lapse imaging of HUVECs labeled with SPY505 and exposed to LSS for 48h. a. siCTRL; b. siSMAD4; c. siALK1. Frame interval – 7.5 minutes.
Flow direction
