## supplementary movie 2 for "Gene-specific endothelial programs drive AVM pathogenesis in SMAD4 and ALK1 loss-of-function"

### Slide 1
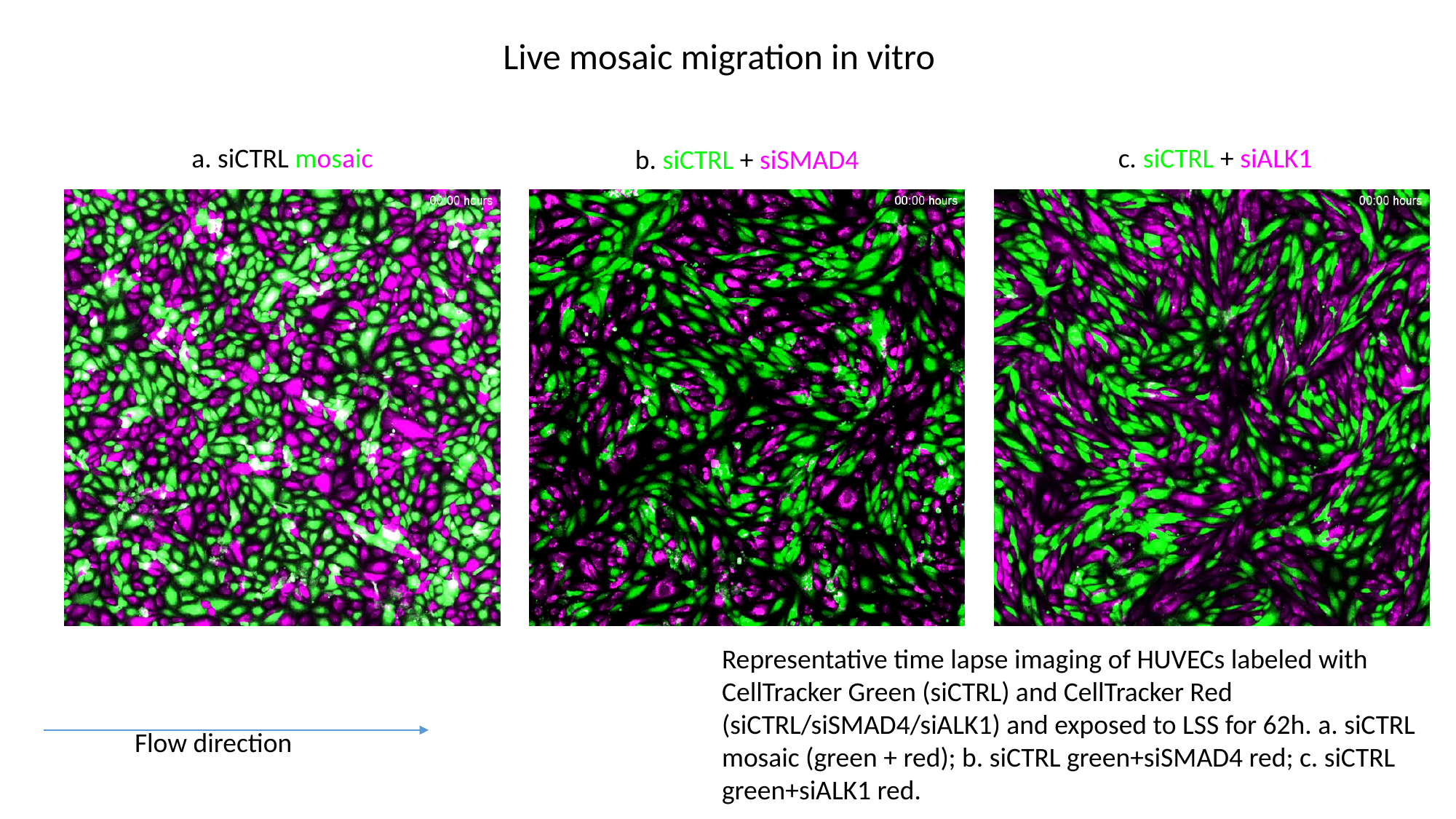

#### Live mosaic migration in vitro
a. siCTRL mosaic
c. siCTRL + siALK1
b. siCTRL + siSMAD4
Representative time lapse imaging of HUVECs labeled with CellTracker Green (siCTRL) and CellTracker Red (siCTRL/siSMAD4/siALK1) and exposed to LSS for 62h. a. siCTRL mosaic (green + red); b. siCTRL green+siSMAD4 red; c. siCTRL green+siALK1 red.
Flow direction
